# Comprehensive Assessment of Smoking and Sex Related Effects in Publicly Available Gene Expression Data

**DOI:** 10.1101/2021.09.27.461968

**Authors:** Emily Flynn, Annie Chang, Bridget M. Nugent, Russ Altman

## Abstract

Smoking greatly reduces life expectancy in both men and women, but with different patterns of morbidity. After adjusting for smoking history, women have higher risk of respiratory effects and diabetes from smoking, while men show greater mortality from smoking-related cancers. While many smoking-related sex differences have been documented, the underlying molecular mechanisms are not well understood. To date, identification of sex differences in response to smoking has been limited to a small number of studies and the resulting smoking-related effects require further validation. Publicly available gene expression data present a unique opportunity to examine molecular-level sex and smoking effects across many tissues and studies. We performed a systematic search to identify smoking-related studies from healthy tissue samples and found 31 separate studies as well as an additional group of overlapping studies that in total span 2,177 samples and 12 tissues. These samples and studies were overall male-biased. In smoking, while effects appeared to be somewhat tissue-specific and largely autosomal, we identified a small number of genes that were consistently differentially expressed across tissues, including *AHRR* and *GZMH*. We also identified one gene, *AKR1C3*, encoding an aldo-keto reductase, which showed strong opposite direction, smoking-related effects in blood and airway epithelium, with higher expression in airway epithelium and lower expression in blood of smokers versus non-smokers. By contrast, at similar significance thresholds, sex-related effects were entirely sex chromosomal and consistent across tissues, providing evidence of stronger effects of smoking than sex on autosomal expression. Due to sample size limitations, we only examined interaction effects in the largest study, where we identified 30 genes with sex differential effects in response to smoking, only one of which, *CAPN9*, replicated in a held-out analysis. Overall these results present a comprehensive analysis of smoking-related effects across tissues and an initial examination of sex differential smoking effects in public gene expression data.

## INTRODUCTION

In some areas of biomedical research, females are underrepresented and sex is still routinely left out of analyses, potentially leading to serious health consequences (Tannenbaum, Day, and Matera Alliance 2017). Many sex and gender differences have been reported in both smoking behaviors and health-related effects of smoking. Smoking is a major cause of premature death, and in the U.S. is estimated to cause more than 480,000 deaths annually (Centers for Disease Control, 2020). After adjusting for smoking history, women have been shown to have increased risk of respiratory symptoms (Langhammer et al. 2000), type 2 diabetes (Will et al. 2001), and lung cancer (Risch et al. 1993). Female smokers also are reported to be 50% more likely to develop COPD than male smokers (Barnes 2016). Despite a higher incidence of smoking-related cancers in females, males have higher mortality from these cancers (Visbal et al. 2004) even though smoking shows a stronger effect on female patient survival (Allen, Oncken, and Hatsukami 2014). However, the biology underlying these differences is not well understood. Improved understanding of the molecular mechanisms behind these smoking-related differences can aid the development of biomarkers and treatments for smoking-related diseases, and may serve as a framework for examining sex differences in other chronic diseases and drug exposures.

Gene expression data provide a unique opportunity to examine molecular level sex differences and dynamic biological responses to smoking. Comprehensive analyses of sex differentially expressed (DE) genes both across (Gershoni and Pietrokovski 2017; Mayne et al. 2016; Oliva et al. 2020) and within individual tissues (e.g. liver (Zhang et al. 2011), blood (Bongen et al. 2019), brain (Trabzuni et al. 2013)) have found hundreds of sex differentially expressed (DE) genes. Additionally, multiple methods (Buckberry et al. 2014; Ellis et al. 2018; Giles et al. 2017; Toker, Feng, and Pavlidis 2016; Flynn, Chang, and Altman 2021) have been developed for inferring sex labels from gene expression data, leveraging the highly sexually dimorphic expression of X and Y chromosome genes. Smoking status also has a substantial impact on gene expression: previous studies have identified hundreds of DE genes between smokers and non-smokers in blood (Charlesworth et al. 2010; Na et al. 2015; Huan et al. 2016), airway epithelium (Chen Xi Yang et al. 2019; Boelens et al. 2009), lung (Landi et al. 2008; He et al. 2018), and other tissues (Port et al. 2004; Na et al. 2015; Tsai et al. 2018). Researchers have found that many of these effects replicate across studies (Huan et al. 2016; Silva and Kamens 2021), and gene signatures predicting smoking status have been identified for blood (Martin et al. 2015; Beineke et al. 2012) and lung tissue (Landi et al. 2008; Bossé et al. 2012).

The impacts of sex and smoking on gene expression vary greatly throughout the body. In the case of sex, the majority of sex-differentially expressed autosomal genes have small, tissue-specific effects, while sex-chromosomal genes generally show consistent differential expression across tissues (Gershoni and Pietrokovski 2017; Mayne et al. 2016; Oliva et al. 2020). By contrast, the tissue-specificity of smoking-related differential expression is less fully characterized. Several analyses have examined effects across tissues, but they focus on cancer (Alexandrov et al. 2016; Desrichard et al. 2018; Alisoltani et al., n.d) and may not extend to healthy tissues.

Characterizing smoking-induced gene expression changes across tissues helps not only with understanding the etiologies of smoking-related cancers, but also may allow for less invasive avenues for sampling. For instance, a blood sample or nasal swab could be used instead of a bronchial brushing or lung biopsy if tissues show substantial overlap in expression. Two studies examined a combination of bronchial epithelium and other epithelial tissues (nasal or nasal and buccal respectively), and found that while there was overlap between smoking-associated DE genes, the majority of DE genes were different between the tissues (Sridhar et al. 2008; Imkamp et al. 2018). Outside of these epithelial tissues, researchers have found less overlap. Morrow and colleagues (Morrow et al. 2019) demonstrated that across airway epithelium, alveolar macrophages, and peripheral blood, samples largely clustered by tissue and there were no shared DE genes; however, there was some overlap in pathway enrichment. Further work is thus required to comprehensively compare the overlap of smoking related effects across a larger number of tissues and studies.

While many studies have examined how smoking and sex individually affect gene expression, to our knowledge, no studies have compared their relative impacts on expression and only a few have identified genes with sex-differential responses to smoking. Consideration and comparison of major drivers of variation is important in biological analysis, and sex differences are often understudied and overemphasized drivers (Patsopoulos, Tatsioni, and Ioannidis 2007). Some sex-related effects may not have clear clinical relevance, so comparison and evaluation of the relative impact of sex-related effects to other drivers of variation (such as smoking and disease states) may shed light on how these factors contribute to health and disease.

In the case of sex-differential smoking effects (also known as sex-by-smoking interaction effects), Yang and colleagues (Chen Xi Yang et al. 2019) identified over 2,500 genes with sex-specific responses to smoking in airway epithelium using data from 211 samples across 16 overlapping studies. In blood, using data from 48 samples, Paul and Amundson (Paul and Amundson 2014) identified 80 genes with sex-differential smoking effects, many of which were associated with female sex hormone receptors (e.g. estrogen and progesterone), and Chatziioannou et al. (Chatziioannou et al. 2017) identified 26 genes with sex-differential effects in 344 blood samples. Identifying and replicating interaction effects is challenging: they are generally very small and require large sample sizes for identification. Across all 3 studies, there is limited overlap of identified genes, which is possibly due to tissue specificity, but further examination of these sex-differential smoking effects is required.

Here, we leverage publicly available gene expression data to examine smoking and sex-related effects at scale and across multiple tissues to identify consistent, reproducible effects. We first perform a systematic search to identify smoking related studies, and then assess sex bias present in these studies. Next, across studies and tissues identified, we compare smoking and sex-related effects and assess the extent to which these effects are shared vs. tissue-specific. Following this, we perform an expanded re-analysis of an airway epithelium dataset to identify smoking, sex, and sex-differential smoking effects. Finally, we attempt to replicate identified sex-differential smoking effects using the largest of our identified studies.

## METHODS

### 1. Identification of smoking-related datasets

#### 1-1 Study search strategy

We identified smoking-related microarray datasets by searching for mentions of the words “smoking/smoker/smoke”, “nicotine”, “tobacco”, or “cigarette” within study and sample metadata. We used a multi-pronged approach to identify smoking-related studies, examining studies from GEO (Edgar, Domrachev, and Lash 2002) and ArrayExpress (Brazma et al. 2003) separately. We used GEOmetadb (Zhu et al. 2008) (downloaded 11/8/2020) to identify GEO human studies and samples that mention a smoking-related term in the metadata. We restricted our sample search to single channel arrays containing either total or polyA RNA samples. We searched for mentions in the “title”, “summary”, or “overall_design” study fields and in the sample “title”, “source_name_ch1”, “treatment_protocol_ch1”, “description”, and “characteristics_ch1” fields. ArrayExpress is the European analog of GEO and contains a large number of expression studies. We searched for mentions of the smoking-related terms in the ArrayExpress browser and downloaded the resulting human studies, filtering for “RNA-seq” and “transcription profiling by array” and removing miRNA platforms. We combined the results of these two searches and removed studies with less than 10 samples.

#### 1-2 Manual Annotation and Filtering

Based on the study title, abstract, and description, studies were manually annotated with tissue type and assigned to one of the following categories:

*1.* **Smokers vs non-smokers or smoking history provided (and at least 1 smoker and 1 non-smoker)**
*2. Treated cells exposed to smoke component*
*3. All smokers (including current vs former)*
*4. All non-smokers*
*5. Not relevant (including cells with other exposures) or no smoking history provided*

#### 1-3 Normalization and extraction of covariate data

For smoking history studies, we extracted phenotypic data on *sex, age, race/ethnicity/ancestry, BMI, and pack years,* where available. *Tissue* annotations were manually assigned. We additionally extracted terms related to disease state (e.g. COPD, cancer) if they were present. Where present, the race/ethnicity/ancestry labels had highly variable annotations across studies. We made efforts to normalize these labels into a combined race/ethnicity/ancestry category, which included African, European, and Asian ancestries, and Hispanic/Latino ethnicity.

### 2. Assessment of sex bias

Our previously developed method for logistic regression-based models for sex labeling (Flynn, Chang, and Altman 2021) were trained on normalized data from the refine-bio database (Greene et al. n.d). This database consists of over 14,000 human studies from GEO, ArrayExpress, and SRA; however, it is not complete. Of the 176 smoking history studies, 139 were contained in refine-bio. For application at scale, we restricted our assessment of sex bias to these 139 studies. As in (Flynn, Chang, and Altman 2021), we grouped studies into the following categories based on the sample sex labels:

*1. Unlabeled:* studies with either less than half of their samples labeled (for studies with up to 60 samples) or less than 30 samples labeled (for studies with more than 60 samples)
*2. Male-only:* all male labels
*3. Female-only:* all female labels
*4. Mostly-male:* >80% of labeled samples are male
*5. Mostly-female:* >80% of labeled samples are female
*6. Mixed sex:* ≤80% of labeled samples belong to either sex

To calculate the fraction of studies that are mixed sex or single sex, we exclude the “mostly” and unlabeled studies from the total and calculate the ratio:

~~~
  frac_mixed_sex = n_mixed_sex / (n_female_only + n_male_only + n_mixed_sex)
  frac_single_sex = (n_female_only + n_male_only)/(n_female_only + n_male_only +
n_mixed_sex)
~~~

### 3. Identification and processing of studies for follow up analysis

#### 3-1 Creation of an Airway Epithelium dataset

There were a large number of airway epithelium studies (n=35) from the same lab and platform (GPL570), many of which contained some of the same sets of samples (Carolan et al. 2006; Harvey et al. 2007; Ammous et al. 2008; Carolan et al. 2008; Tilley et al. 2009; Vanni et al. 2009; Hübner et al. 2009; Raman et al. 2009; Carolan et al. 2009; Leopold et al. 2009; Turetz et al. 2009; Dvorak et al. 2011; Strulovici-Barel et al. 2010; R. Wang et al. 2010; Shaykhiev et al. 2011; Marcus W. Butler et al. 2011; M. W. Butler et al. 2011; R. Wang et al. 2011; Tilley et al. 2011; Hackett et al. 2012; R. Wang et al. 2012; Buro-Auriemma et al. 2013; Shaykhiev et al. 2013; Gao et al. 2014; Hessel et al. 2014; Walters et al. 2014; Tilley et al. 2016; Zhou et al. 2016; J. Yang et al. 2017; G. Wang et al. 2017) (see **Supplementary Table 1** for a list of study accessions and titles). We aggregated these samples into a *Grouped Airway Epithelium (Grouped AE)* dataset. Many of the samples contain covariate information related to age, race/ethnicity and pack-years (see Table 1A). The dataset contains both large and small airway epithelium samples, which largely cluster together in principal components space (see **Supplementary Figure S6A**).

**Table 1.** Smoking and sex breakdown of airway epithelium data

|  |  | Smokers |  |  | Non-smokers |  |  |
| --- | --- | --- | --- | --- | --- | --- | --- |
|  |  | total | female* | male | total | female | male |
| **n** |  | 273 | 73 | 200 | 171 | 61 | 110 |
| **age<sup>▽</sup>** | **mean ± sd** | 42.6±7.4 | 41.3±8.9 | 43.1±6.8 | 40.3±10.2 | 37.9±11.3 | 41.6±9.4 |
|  | **missing** | 79 | 19 | 60 | 30 | 13 | 17 |
| **race<sup>+</sup>** | **Asian** | 0 | 0 | 0 | 4 | 4 | 0 |
|  | **Black** | 119 | 33 | 86 | 67 | 20 | 47 |
|  | **Black, Hispanic** | 0 | 0 | 0 | 2 | 2 | 0 |
|  | **Hispanic** | 32 | 7 | 25 | 20 | 8 | 12 |
|  | **White** | 45 | 14 | 31 | 50 | 14 | 36 |
|  | **missing** | 77 | 19 | 58 | 28 | 13 | 15 |
| **pack years<sup>◊</sup>** | **mean ± sd** | 27.6±16.8 | 27.1±16.4 | 27.7±17.1 | -- | -- | -- |
|  | **missing** | 81 | 20 | 61 | -- | -- | -- |

\*Sex is not significantly associated with smoking p = 0.059 (chi-squared test)
▽Age is associated with smoking status (p = 0.02) and sex is also associated with age (p=0.01). Missingness of age associated with smoking (p=0.009) but not sex (p=0.92).
+Race-ethnicity is significantly associated with smoking status (chisq p = 0.03, removed categories with less than 5 counts total) but not sex (p=0.99). Missingness of race-ethnicity associated with smoking status (p=0.006) but not sex (p=1)
◊Pack-years is not associated with sex (p=0.8) (t-test) and missingness of pack-years is not associated with sex (p=0.29) or race (p=0.08)

For processing, we first filtered to remove samples from subjects with COPD or asthma, and for subjects with repeated measures, we used the first sample from the subject. We then downloaded the raw expression data from GEO and used the R package affy (Gautier et al. 2004) to load, normalize, and RMA transform the data. Many of the samples were direct duplicates across studies. For these samples, we combined their metadata, which exactly matched for sex and race with the exception of one sample which we excluded. Three samples contained different but nearby ages or pack-years; we took the average of the two values. We also grouped by study participant ID (or “DGM” id in the metadata) and removed repeated samples with the same participant ID.

Prior to covariate imputation and modeling, we grouped together categorical values with small n. For race-ethnicity labels, we assigned samples in race-ethnicity groups with less than 5 counts to “other race-ethnicity” for modeling purposes. Sample submission date correlated with expression, but contains 22 variables, many with small counts. For date groups with less than 10 counts, we assigned the samples to the nearest submission date with more than 10 counts, resulting in 10 total submission date categorical variables. We chose to do this (rather than assigning all samples with small numbers of counts to an “other date” category) because samples appear to cluster together over time in PC space (see **Supplementary Figure S6B and C** for before and after date collapsing).

#### 3-2 Systematic Search for Smoking Studies across Tissues

After removing the overlapping airway epithelium datasets, we focused on identifying studies using healthy tissues from at least 5 never smokers and current smokers at the time of sample selection. To do so, we downloaded the sample-level metadata for these studies in order to determine if there were sufficient samples. We included healthy tumor adjacent tissue from individuals with cancers, but excluded samples from individuals with COPD or other annotated diseases. We also removed studies from single sex tissues (prostate) or associated with pregnancy (placenta, umbilical cord). For studies with repeated samples from the same subject, we include only the first sample. We also did not include “ever” smokers unless additional information was present indicating that they were still smoking.

For quality control, we inferred sample sex labels for candidate studies. While our penalized logistic regression model performs well at scale, clustering based methods are better for examining large mixed sex studies because they allow for visualization and examination of within-study clustering. Where expression levels for *XIST, RPS4Y1*, and *KDM5D* were available, we applied the Toker method (Toker, Feng, and Pavlidis 2016), otherwise we used massiR (Buckberry et al. 2014), which clusters based on the expression of Y chromosome genes. We manually checked each study to ensure clear separation and excluded six studies, and excluded mislabeled samples and studies without clear sex separation.

#### 3-3 Processing of Small Expression Studies

MetaIntegrator (Haynes et al. 2017) was used to download the data as processed by the authors. MetaIntegrator performed log-transformation and quantile normalization if these steps were not already taken.

### 4. Variance Decomposition

We sought to examine the fraction of variance in each dataset associated with smoking and the sex-by-smoking interaction effects. To do this, we used principal variance components analysis (PVCA). Briefly, this method first performs PCA and then identifies the cumulative fraction of the variance explained by each of the covariates in a model across the first n PCs, where n is chosen based on the number of PCs that explain a cutoff fraction of the total variance. We used 0.8 for the cutoff fraction, but obtained similar results across a range of cutoffs (0.4-0.9). The R package variancePartition (Hoffman and Schadt 2016) was used to calculate the variance fractions.

We ran PVCA with two models:

1. baseline model: PC_i_ ∼ sex + smoking + C
2. interaction model: PC_i_ ∼ sex + smoking + sex*smoking + C

where C is the set of additional covariates, and PC_i_ is the ith PC.

The cumulative variance for covariate j is given by ∑ (X_ij_ * v_i_) where X_ij_ is the fraction of the variance in PC_i_ explained by covariate j and v_i_ is the fraction of the total variance in the expression data explained by PC_i_.

### 5. Differential expression analysis

#### 5-1 Differential expression model

We performed differential expression analysis separately on each of the small datasets and the grouped airway epithelium dataset. The R package limma (Ritchie et al. 2015) was used for differential expression analysis, with the following model:

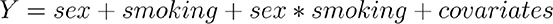

Sex and sex*smoking covariates were excluded from single-sex datasets. We used the cutoffs FDR <0.05 and absolute effect size log fold change of ≥0.3 for identifying differentially expressed (DE) genes.

#### 5-2 Summarizing probes to genes

Because the studies spanned a variety of platforms, identification of DE genes and comparison across studies was performed at the gene level. Probes were mapped to HGNC gene symbols using the appropriate Bioconductor package (hgu133plus2.db, hgu219.db, hgu133a.db, hgu133a2.db, hugene10sttranscriptcluster.db) for five platforms. For the remaining 7 platforms, the probe-to-gene mapping was downloaded directly from GEO.

For meta-analysis, following model fitting at the probe-level, we used fixed effects inverse variance meta-analysis to summarize effect sizes to genes, as implemented in the R package meta (Schwarzer, Carpenter, and Rücker 2015).

Due to the lack of ground truth, we chose to drop out portions of a dataset and apply these methods, where the “true” genes were the DE genes from the full dataset (where DE genes are the set of genes to which all DE probes mapped). We used the Grouped AE dataset as the full dataset, and smoking as the covariate examined. We examined the precision and recall of the three methods at two FDR cutoffs (< 0.01 and < 0.05) and across varying dropout fractions (0.3-0.9), with fifteen random dropouts per fraction (see **Supplementary Figure S9** for the results). For this analysis, we wanted to be conservative in our estimates, and as a result, chose to use meta-analysis for summarization.

#### 5-3 Assessment of replication and overlap

Genes identified in the Grouped AE dataset were replicated using the dataset GSE7895, which was selected for validation because it was the largest airway epithelium dataset present in the set of smaller studies. We identified lists of *replicated genes*, which we define as the subset of DE genes from the discovery that have a p-value < 0.05 in the validation and effect sizes in the same direction in the discovery and validation sets. We also examined the correlation between the effect sizes of the DE genes.

For examining overlapping genes between 2 studies (rather than replication), we use the union of the DE genes (FDR < 0.05, logFC ≥0.3), resulting in *n overlapping genes*. We identify *overlapping significant genes* as genes that have effect sizes in the same direction and p-value < 0.05/*ngenes* in both studies where *ngenes* is the number of overlapping genes. In order to examine the similarities between 2 studies related to their association with the variable of interest (smoking, sex), we examined the correlation of the effect sizes. We used a permissive cutoff for genes included (FDR < 0.10 in either study) and, if there were at least 30 genes remaining, we calculated the correlation coefficient across genes for mean effect sizes weighted by their standard deviations. We chose to use a weighted correlation coefficient in order to be less sensitive to the FDR cutoff, while ensuring that genes with smaller standard errors are weighted more highly.

#### 5-4 Examining tissue specificity

We used to examine tissue specificity of particular genes and compare the tissue-specificity between smoking- and sex-related analyses (Yanai et al. 2005). This metric was designed for examining tissue-specificity of expression of a particular gene and results in a number 0 to 1 where 0 is ubiquitously expressed and 1 is tissue-specific. We extend this to examine tissue-specificity of differential expression by inputting the absolute log fold-change values instead of the log expression intensity to obtain the tissue-specificity of differential expression. The formula for τ is given below:

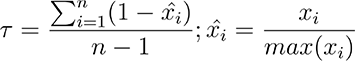

Where *n* is the number of tissues and we define *x_i_* as the median log-fold-change in tissue *i*. Importantly this does not distinguish between opposite direction effects, so it is important to also examine their presence.

### 6. Between- and within-tissue meta-analyses

We performed random effects meta-analysis using the DerSimonian-Laird estimator, first across studies and tissues, and then for blood and airway epithelium studies separately, examining both smoking and sex-related effects. The Grouped AE study was not included in the meta-analysis because it is substantially larger than the other datasets and as a result may have a strong impact on the results. We selected 4 validation studies: *GSE7895* (airway epithelium), *GSE27002* (alveolar macrophages)*, GSE21862* (PBMCs), and *E-MTAB-5279* (whole blood).

We included 6 whole blood and two PBMC studies in the blood meta-analysis. The B cell study was excluded because it represents a specific cell type in blood, while the others are a mixture (meta-analysis of all blood studies including the B cell study also shows similar results). For the airway epithelium meta-analysis, we included four airway epithelium studies and added the trachea epithelium study because trachea is an airway tissue and overlaps in PC space (we expect this may reflect differences in terminology), and the expression was highly correlated.

We performed the smoking-related meta-analysis for genes present in at least 15 of the 27 studies. For the sex-related meta-analysis, we selected a lower cutoff for number of studies (n=10 out of 24) because of the large number of missing sex chromosome probes. Finally, for blood and airway meta-analyses, we filtered for at least 5 blood and 4 airway studies respectively.

For validation, we considered a gene validated in a particular study if the gene’s effect size is in the same direction and has a p-value < 0.05 / (number of genes).

### 7. Sample size calculation for interaction effects

We examined the sample size required to detect an interaction effect in an expression dataset in the case where we have two binary covariates (smoking, sex) and under the assumption that the data is balanced. We used the R package ssize (Warnes et al. 2020) with a power of 0.80 and FDR of 0.05. We assumed uniform standard deviations of probes, and used a value of 0.6 based on the mean empirical standard deviation of probes across datasets included. We then examined the sample size required for detecting absolute log effect sizes in the range of 0.1 to 0.6, assuming 90%, 95%, and 99% of genes were not differentially expressed (see **Supplementary Figure S10**).

## RESULTS

### 1. Systematic search for smoking-related studies

We performed a systematic search of human gene expression studies in GEO and ArrayExpress to identify studies that have smoking-related information (see **Supplementary Figure S1** for a diagram showing the systematic search approach). We searched both sample and study metadata and identified 530 studies (spanning 63,772 samples) that contained a smoking-related mention. We manually annotated the studies to identify the subset that have smoking history information (n=176 studies).

To examine effects across tissues, we identified the subset of smoking history studies that contain samples from at least 5 healthy smokers and non-smokers (see Table 1B for the list and their sample breakdown). Thirty-five studies in airway epithelium were from the same lab, using the same microarray platform, and had many overlapping samples. We combined all of these into a single larger study (further described as *Grouped Airway Epithelium or Grouped AE*), which contained 444 samples after deduplication (see Table 1A**, Methods 3-3**). The additional airway epithelium studies are distinguished from the *Grouped AE* study in that they are either from another lab and/or on a different microarray platform.

**Table 1B.** Sex breakdown of smaller studies organized by tissue. The number of females in each category is included in parentheses.

| tissue | study | title (citation where available) | platform | smokers | non-smokers | sex label mismatch | additional phenotypes |
| --- | --- | --- | --- | --- | --- | --- | --- |
| airway epithelium | GSE14633 | Gene expression from bronchial epithelial cell samples of current and never smokers.(Schembri et al. 2009) | GPL5175 | 11 (3) | 11 (7) | 0 | race; pack years |
| airway epithelium | GSE19027 | Antioxidant response gene expression in the bronchial airway epithelial cells of smokers at risk for lung cancer (X. Wang et al. 2010) | GPL96 | 22 (1) | 7 (2) | 2 | age; race; pack years |
| airway epithelium | GSE4302 | Genome-wide profiling of airway epithelial cells in asthmatics, smokers and healthy controls (Woodruff et al. 2007) | GPL570 | 15 (2) | 28 (16) | no metadata | NA |
| airway epithelium | GSE5056 | Airway epithelium, large airways, phenotypically normal smokers vs non-smokers, MAS5 (HuGeneFL) (Carolan et al. 2006) | GPL80 | 26 (8) | 18 (4) | 0 | age; race; pack years |
| airway epithelium | GSE7895 | Reversible and permanent effects of tobacco smoke exposure on airway epithelial gene expression (Beane et al. 2007) | GPL96 | 52 (10) | 21 (5) | 0 | age; pack years |
| alveolar macrophages | GSE13896 | Smoking-dependent reprogramming of alveolar macrophage polarization: implication for pathogenesis of | GPL570 | 50 (6) | 43 (10) | 4 | age; race; pack years |
|  |  | COPD (Shaykhiev et al. 2009) |  |  |  |  |  |
| alveolar macrophages | GSE2125 | Isolated alveolar macrophages (Woodruff et al. 2005) | GPL570 | 13 (2) | 15 (10) | no metadata | NA |
| alveolar macrophages | GSE27002 | Chronic cigarette smoke exposure results in coordinated methylation and gene expression changes in human alveolar macrophages (R. A. Philibert et al. 2012) | GPL5175 | 13 (4) | 10 (5) | no metadata | NA |
| blood - b cells | GSE18723 | Gene expression circulating B lymphocytes for smoking females (Pan et al. 2010) | GPL96 | 38 (38) | 40 (40) | all female | menopause |
| blood - pbmcs | GSE21862 | Gene expression on 144 arrays representing 125 workers exposed to a range of benzene exposures (McHale et al. 2011) | GPL6104 | 9 (1) | 33 (24) | 0 | age; subject_id; batch (chip id) |
| blood - pbmcs | GSE42057 | Peripheral blood mononuclear cell gene expression in chronic obstructive pulmonary disease (Bahr et al. 2013) | GPL570 | 13 (7) | 27 (13) | 0 | age; pack_years; fev1; bmi; activity |
| blood - pbmcs | GSE87072 | Gene expression profiles from PBMCs collected from chronic smokers and moist snuff consumers (Arimilli et al. 2017) | GPL570 | 40 (0) | 40 (0) | all male | age |
| blood - whole | E-MTAB-5278 | Transcription profiling of blood from smokers (with or without COPD), non-smokers and former smokers to identify gene expression signature for cigarette smoke exposure response (Martin et al. 2015) | GPL570 | 56 (23) | 56 (23) | 4 | race; age |
| blood - whole | E-MTAB-5279 | Transcription profiling of blood from smokers, non-smokers and former smokers to identify gene expression signature for cigarette smoke exposure response (Martin et al. 2015) | GPL570 | 27 (12) | 28 (13) | 0 | race; age |
| blood - whole | GSE20189 | A gene expression signature from peripheral whole blood for stage I lung adenocarcinoma (Rotunno et al. 2011) | GPL571 | 27 (14) | 21 (11) | no metadata | NA |
| blood - whole | GSE20681 | Whole blood cell gene expression profiling in patients with coronary artery disease from the PREDICT trial (Elashoff et al. 2011) | GPL4133 | 14 (3) | 48 (16) | 0 | age |
| blood - whole | GSE23323 | Transcriptomics in response to cigarette smoking in humans | GPL6480 | 22 (10) | 22 (10) | no metadata | NA |
| blood - whole | GSE23515 | Radiation responses in peripheral white blood cells of smokers and non-smokers (Paul and Amundson 2011) | GPL6480 | 12 (6) | 12 (6) | 0 | age |
| blood - whole | GSE56768 | Whole blood and isolated blood cell transcriptomics in COPD | GPL570 | 39 (19) | 5 (3) | no metadata | NA |
| brain - prefrontal cortex | GSE30272 | Temporal dynamics and genetic control of transcription in the human prefrontal cortex | GPL4611 | 56 (23) | 166 (52) | 0 | race; age; alcohol; postmortem |
|  |  | (Colantuoni et al. 2011) |  |  |  |  | interval;<br>batch |
| buccal<br>mucosa | GSE16149 | Examining smoking-induced differential gene expression changes in buccal mucosa (Kupfer et al. 2010) | GPL570 | 9 (9) | 9 (9) | all female | NA |
| buccal<br>mucosa | GSE17913 | Effects of cigarette smoke on the human oral mucosal transcriptome (Boyle et al. 2010) | GPL570 | 35 (16) | 33 (16) | 9 | NA |
| kidney | GSE46699 | Smoking and obesity related molecular alterations in clear cell renal cell carcinoma (Eckel-Passow et al. 2014) | GPL570 | 21 (7) | 37 (22) | no<br>metadata | obesity |
| liver | GSE32504 | Identification of expression quantitative trait loci (eQTL) in human liver (Schröder et al. 2013) | GPL13376 | 28 (12) | 115 (64) | 2* | race; age;<br>alcohol;<br>medication |
| lung | GSE103174 | Expression data from lung tissue in mild-moderate COPD | GPL13667 | 5 (2) | 10 (9) | 1 | age; bmi;<br>pack years;<br>fev1; batch;<br>cell types |
| lung | GSE31210 | Gene expression data for pathological stage I-II lung adenocarcinomas (Okayama et al. 2012) | GPL570 | 11 (4) | 7 (4) | 0 | age |
| lung | GSE32539 | Molecular phenotyping of the idiopathic interstitial pneumonias identifies two subtypes of idiopathic pulmonary fibrosis (I. V. Yang et al. | GPL6244 | 21 (11) | 20 (5) | 1 | age; rin;<br>pack years;<br>batch<br>(aliquot) |
|  |  | 2013) |  |  |  |  |  |
| nasal epithelium | GSE8987 | Expression data from buccal and nasal epithelium of current and never smokers (Sridhar et al. 2008) | GPL571 | 7 (1) | 8 (2) | no metadata | NA |
| oral cavity | GSE42743 | Oral cavity cancer compared to adjacent "Normal" tissue [validation set] (Lohavanichbutr et al. 2013) | GPL570 | 11 (3) | 6 (2) | 0 | age/dxdate |
| sputum | E-MTAB-3604 | Alterations in the sputum proteome and transcriptome in smokers and early-stage COPD patients (Titz et al. 2015) | GPL570 | 40 (15) | 45 (17) | 0 | race; age; pack years; bmi; fev/fvc |
| trachea epithelium | GSE994 | Effects of cigarette smoke on the human airway epithelial cell transcriptome (Spira et al. 2004) | GPL96 | 31 (7) | 18 (4) | no metadata | NA |

\*For comparison, we used the paper supplement metadata for this study, GEO metadata showed exactly the opposite sex labels.

The remaining 31 studies (1754 samples) are majority blood or blood component (n=11), followed by airway epithelium (n=5), then lung and alveolar macrophages (n=3), and buccal mucosa (n=2), and 1 each of nasal epithelium, tracheal epithelium, oral cavity, sputum, kidney, liver, and brain (prefrontal cortex). While the lower bound was 5 smokers and non-smokers, the range for identified studies was 5 to 166 smokers and 5 to 56 non-smokers (medians = 21 and 22 respectively). Seven studies had significantly more smokers (p= 1.6*10^-13^ to 4.7*10^-2^) while 3 had significantly more non-smokers (p= 3.0*10^-7^ to 5.4*10^-5^).

**Figure 1.**
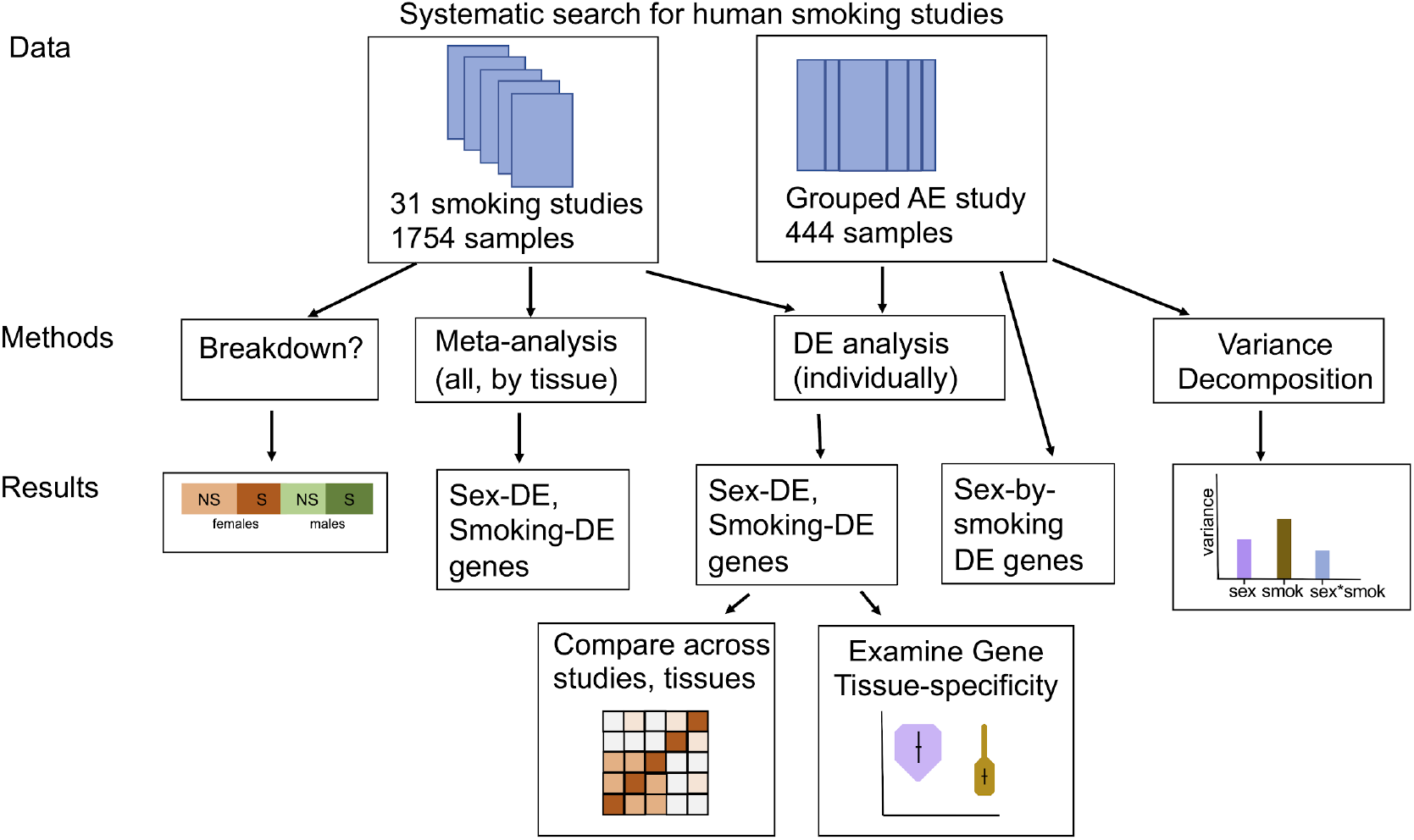
Study schematic. We performed a search for human gene expression studies on smoking. This resulted in a set of 31 separate studies, as well as a group of overlapping airway epithelium (AE) studies we combined into a single grouped study. We examined the sex breakdown in these studies and perform both individual differential expression analyses as well as meta-analyses across studies and tissues in order to identify differentially expressed genes. We used the results of these analyses to compare the effects of smoking and sex across studies and tissues.

### 2. Smoking-related samples are male-biased

We additionally sought to examine sex bias overall in smoking-related studies. We focused on the 139 (out of 176) smoking history studies that were included in the refine-bio database by inferring sex labels from gene expression data using our previously published method (Flynn, Chang, and Altman 2021). For smoking history studies, 34.5% of samples and 38.8% of studies were missing metadata sex labels; this is much lower than seen across all human studies and samples (e.g. 70.7% of human microarray samples are missing sex labels (Flynn, Chang, and Altman 2021)). The higher fraction of sex labels in smoking datasets may be related to the fact that smoking status is included, so sex is additionally likely to be recorded as a covariate.

After inferring sex labels from expression, we found that smoking-related samples are slightly male-biased with 59.1% and 68.1% percent of labeled samples derived from males for smoking history and treated cell studies, respectively. This is in contrast to the overall pattern of human samples which is slightly female-biased (52.1%) but matches the pattern that more men smoke. The majority of smoking history studies are mixed sex (92% of labeled studies). The high fraction of mixed sex studies helps with follow up examination of sex-related effects (see **Supplementary Figure 2** and **Supplementary Table S2** for the sample and study sex breakdowns, respectively).

Of the 31 studies included in our follow up analysis, 9 did not have metadata sex labels and 3 studies were single sex. In addition to the higher proportion of males (59.4%, p < 4*10^-15^), male sex was also significantly associated with smoking status (p < 0.0007, see **Supplementary Figure S3** for the sex and smoking breakdown of these studies). Seven studies contained a total of 23 samples where the inferred sex did not match the metadata sex, corresponding to 1.3% of the samples examined (see Table 1B). The Grouped AE study was a higher fraction male (70%) and contained 2.4% mislabeled samples (see Table 1A**, Supplementary Figure S7**). Sample sex mismatches highlight the potential for mislabeled samples along other dimensions (e.g. smoking status), and were excluded from follow up analysis.

### 3. Smoking effects are largely tissue-specific *and autosomal* but show some consistency across tissues, *while sex-related effects are sex chromosomal and consistent across tissues*

We sought to examine the extent to which smoking-related effects are consistent across the tissues and the studies we examined. First, we performed differential expression analysis within each study across tissues (airway epithelium, lung, kidney, buccal mucosa, etc.) (see **Supplementary Table S3** for a summary of results across studies), and summarized probes to genes with meta-analysis. Four studies showed no differentially expressed (DE) genes related to smoking, while the remaining studies had between 2 and 4357 DE genes, with a median of 31. As expected, larger studies had more DE genes (for smoking: spearman’s =0.36, p = 0.049, sex and sex-smoking n.s.) and more overlap between each other.

Overlap and between-study correlations of smoking-related effects appear to cluster by tissue, with separate clusters of airway epithelium and blood studies (Figure 2A shows the counts of overlapping genes; Figure 2C contains the correlations of top genes between all pairs of studies). For example, Grouped AE showed the highest correlation with other airway epithelium studies ( =0.72, 0.57, and 0.55) and the trachea epithelium study ( = 0.584). By comparison, sex-related effects appear to correlate across studies and tissues (see Figure 2D). We separated out the autosomal (Figure 2E) genes, and found that the strong pattern of shared, consistent sex-related effects is largely limited to the sex chromosomes.

**Figure 2.**
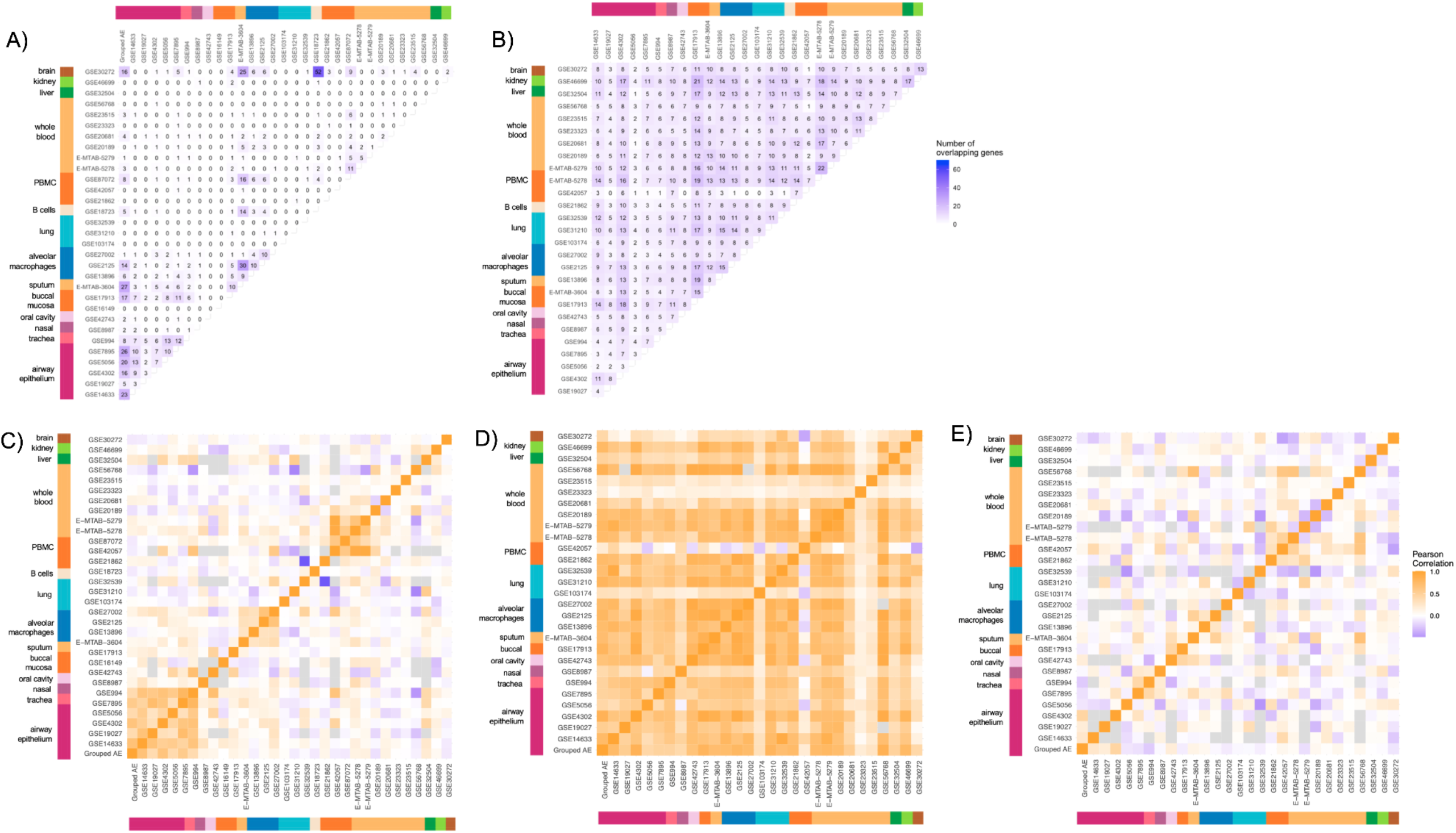
Smoking and sex-related effects across tissues. Heatmaps showing the numbers of overlapping significant genes **(A-B)** and correlation of effect sizes **(C-E)** in each of the studies for smoking (A, C) and sex-related effects (B, D, E). Studies are organized by tissue, as indicated by the color bars on the side. A-B) The number of overlapping genes is shown, with darker purple indicating larger numbers for smoking (A) and sex (B). C-E) Correlation plots colored by weighted Pearson correlation of the effect sizes (weighted by standard error) for top overlapping genes (FDR < 0.1 in either study). Correlations are plotted for smoking-related effects (C) and for sex-related effects separated into sex chromosome (D) and autosomal effects (E). Orange indicates a positive correlation, white indicates no correlation, and purple is a negative correlation. Correlation was not calculated for pairs of studies with less than 30 overlapping genes; these are shown in gray.

While the majority of overlap clustered by tissue, 7 DE genes were present in 5 studies spanning both an airway-related tissue (airway, sputum, oral, buccal, lung, or alveolar) and non-airway tissue (blood, brain, kidney or liver): *LRRN3, MS4A6A, GAPDH, RPLP0, CX3CL1, GPR15,* and *AHRR* (another 7 genes were present in 4 studies with both an airway and non-airway), indicating the presence of some consistent smoking-related effects across tissues (see **Supplementary Table S4A** for full lists of smoking DE genes present in at least two studies).

We also performed a meta-analysis across tissues using 27 out of 31 studies (see **Methods 6**), and identified 7 genes that showed significant smoking-related effects: the expression of *AHRR, CYP1B1, NQO1, LRRN3* were significantly higher and *ELOVL7*, *CCL4*, and *GZMH* were significantly lower in current smokers as compared to non-smokers (see **Supplementary Table S5** for their effect sizes). Figure 3A shows the study-level expression of these 7 genes as well as the pooled estimate. In our analysis, we identified *LLRN3* and *AHRR* as genes that had an effect in both an airway and non-airway tissue. Two genes, *GZMH* and *AHRR,* appear to show relatively consistent effects across tissues, showing consistently lower and higher expression in smokers vs. non-smokers respectively. For the remainder of these genes, the effects appear to be tissue-dependent. *NQO1* shows a strong association with smoking in airway epithelium, while *LRRN3* appears to show a stronger association with smoking in blood (both have higher expression in smokers). *CYP1B1* shows strongest association with smoking in airway epithelium (higher in smokers), while *ELOVL7* and *CCL4* appear to be strongest in alveolar macrophages and sputum (lower in smokers).

**Figure 3.**
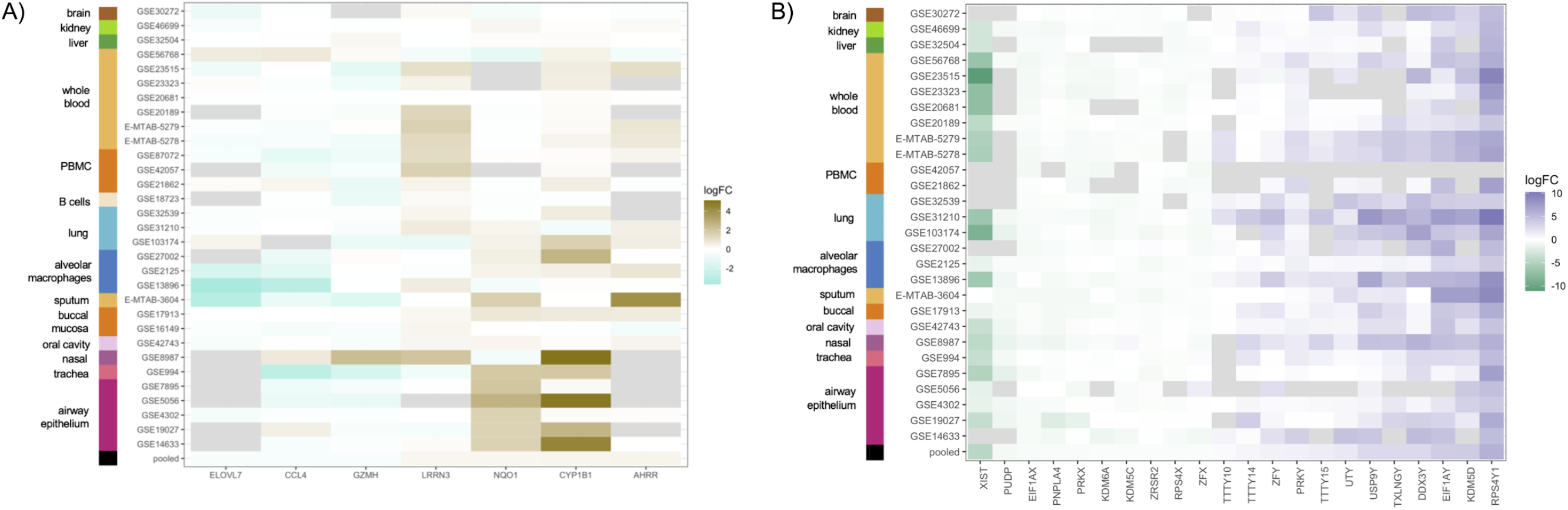
Meta-analysis of differential expression across studies for smoking **(A)** and sex **(B).** Studies are organized by tissue, as indicated by the color bars on the left side of each heatmap. The color of the heatmap tiles show the log-fold change (logFC) of the association between the variable of interest (smoking or sex) and that gene in that specific study: gold is more highly expressed in smokers and turquoise is more highly expressed non-smokers, green is higher in females and purple is higher in males. Gray tiles indicate missing values.

We examined whether these genes were differentially expressed in four held-out validation datasets (*GSE7895* - airway epithelium, *GSE27002* - alveolar macrophages, and *GSE21862* and *E-MTAB-5279* - blood). Four of the smoking-related genes were differentially expressed in the validation datasets, each in one study: *LRRN3* (blood), *AHRR* (blood), *NQO1* (airway epithelium), and *CYP1B1* (alveolar macrophages). Interestingly, *LRRN3* and *NQO1* showed similar tissue-specificity to the discovery dataset.

#### Although some genes showed consistent responses to smoking across tissues, looking within tissues highlights key genes involved in tissue-specific responses

We performed tissue specific meta-analyses for blood and airway epithelium studies. The blood analysis included two PBMC and five whole blood studies, while the airway epithelium analysis included four airway and one trachea epithelium study (see **Supplementary Figure S4** for heatmaps and **Supplementary Table S5** for the lists of genes). At an FDR of 0.05 and effect size cutoff of ≥0.3, the blood meta-analysis identified 19 DE genes, while the airway epithelium analysis identified 66 DE genes. In airway epithelium, 21 out of the 66 DE genes validated in the held-out airway epithelium dataset (*GSE7895*). In blood, only 3 DE genes were replicated (*SH2D1B, KLRF1, AKR1C3*). Only 1 gene, *AKR1C3*, overlapped between the 2 meta-analyses and interestingly, it showed opposite direction effects in the 2 tissues (pooled effect size estimates: logFC=-0.32, p= 2.0*10^-5^ in blood and logFC=1.6, p = 6.2*10^-10^, both validated), as shown in the violin plot in Figure 4.

**Figure 4.**
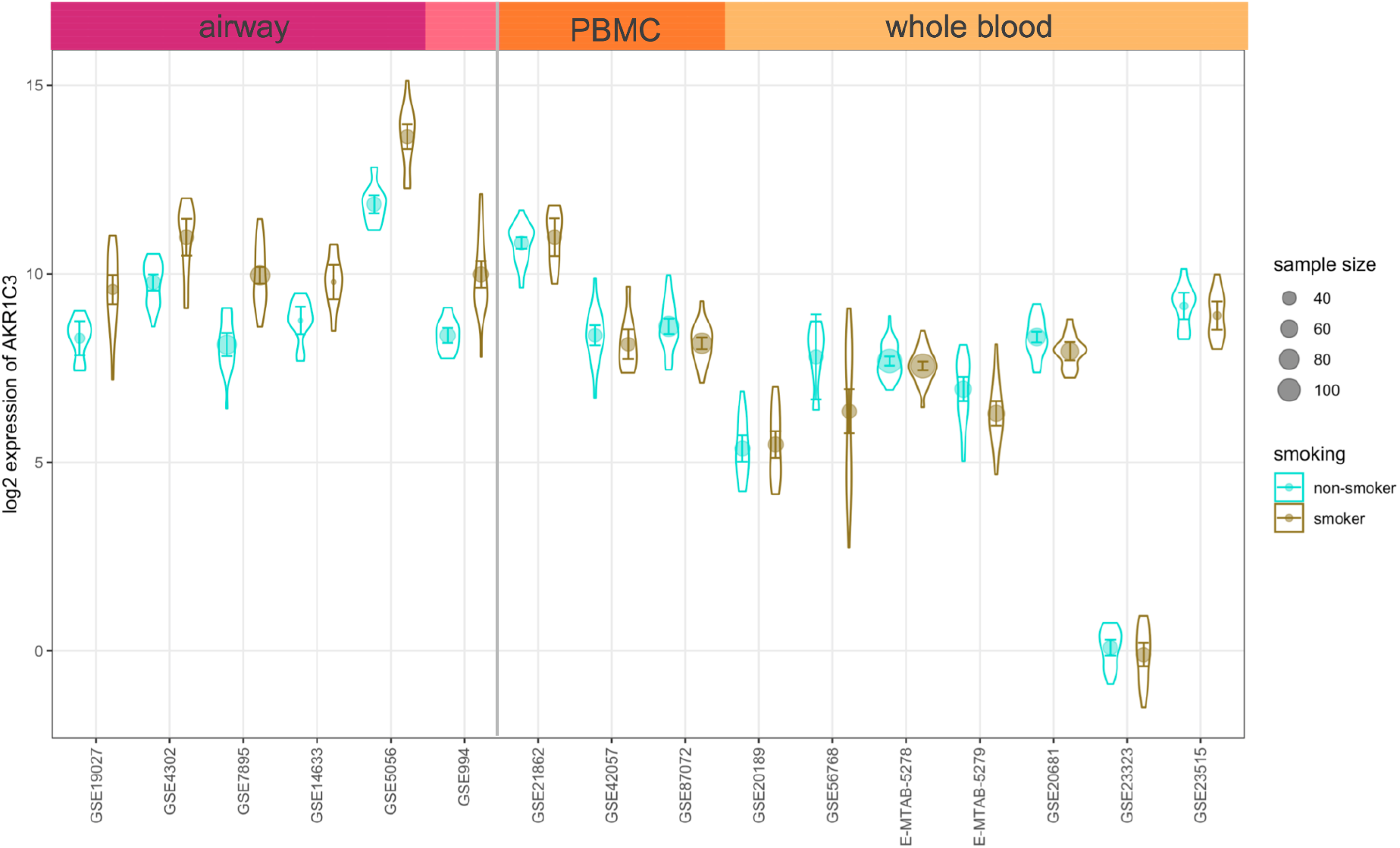
Violin plot showing the distribution of *AKR1C3* levels across smokers (gold) and non-smokers (teal) in airway and blood studies. The mean and 95% confidence interval are included for each study/smoking group, and the size of point corresponds to the overall study sample size.

By contrast, most sex-DE genes were consistent across studies and tissues: forty-five genes were consistently DE in at least three studies (see **Supplementary Table S4B**). Only four of these genes were autosomal (*EIF5B, ACTB, KLF6, LAPTM4B*), and the sex-DE autosomal genes had higher expression in females. Six of the DE sex chromosomal genes were present in 20 or more studies, including *RPS4Y1, EIF1AY, DDX3Y, KDM5D, UTY, USP9Y,* and *XIST*. We additionally saw little evidence of tissue specificity for the sex-related meta-analysis (Figure 3B), which identified 22 X and Y chromosome genes with sex differences in expression: 12 higher in males and 10 higher in females. All but 2 of these genes validated in a held-out dataset, and 11 validated in 2 or more datasets. Tissue-specific, sex differences meta-analyses resulted in 32 genes in blood and 6 in airway epithelium. The majority of these genes were sex chromosomal; however, 15 genes in blood and 1 gene in airway epithelium were autosomal. Overall, 14 blood and 4 airway epithelium genes validated in the held-out datasets; all validated genes were sex chromosomal.

It is important to note that for analysis, we inferred sex labels using the expression of a subset of X and Y chromosome genes (although there are many other X and Y genes that are DE). In addition, when we examined the subset of studies with metadata sex labels (35 studies) and assumed that these labels were correct, we obtained similar patterns of significantly differentially expressed X and Y chromosome genes that were overlapping across studies and tissues.

#### Genes associated with smoking show more tissue specificity than genes with similar effect sizes associated with sex

We examined the subset of DE genes present in at least 3 studies and 2 tissues, and adapted the τ tissue-specificity metric (Yanai et al. 2005) to examine specificity of differential rather than absolute gene expression (see **Methods 5-4**). Across DE genes, smoking-related genes showed significantly more tissue-specificity than sex-related related genes (p= 4.92 * 10^-10^) (Figure 5A for the summary of these effects and **Supplementary Figure S5** to visualize differences at the gene level).

**Figure 5.**
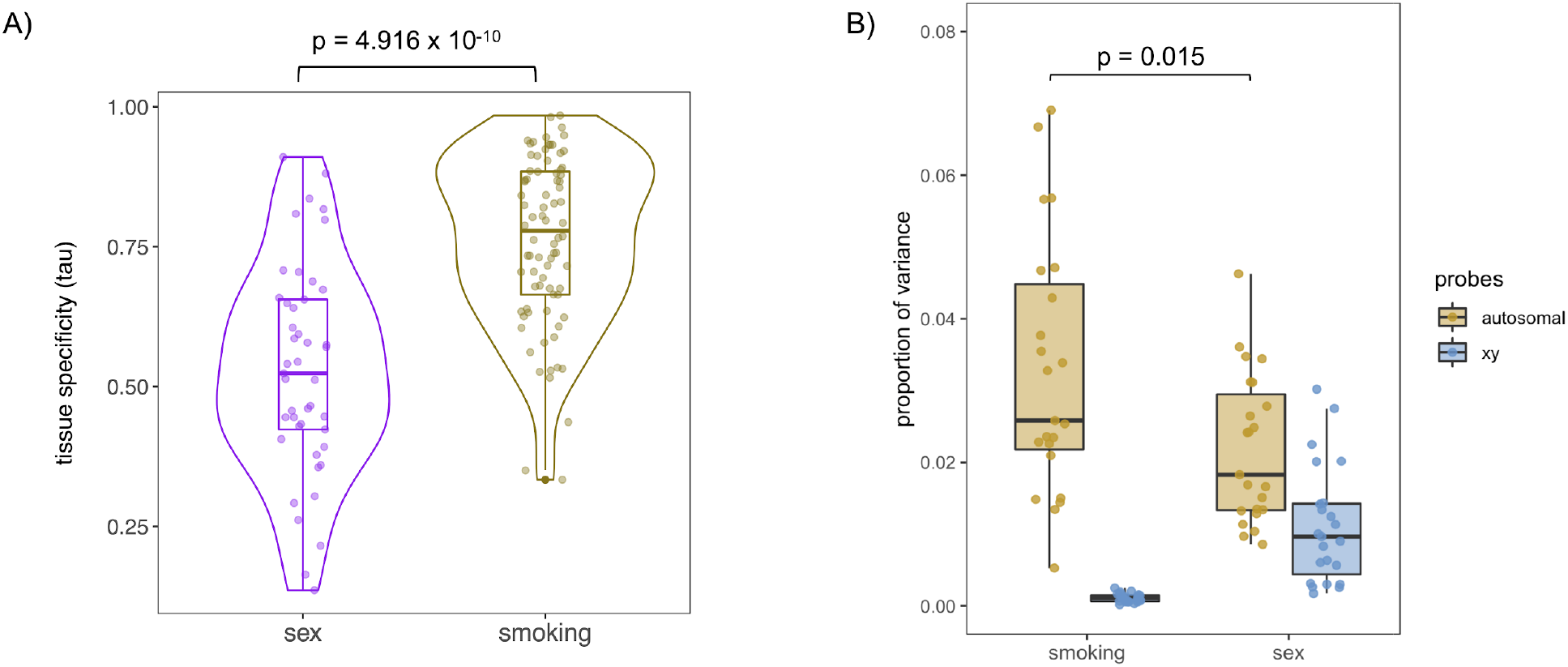
Comparison of sex and smoking effects. **(A)** Smoking-related genes (gold) show higher tissue-specificity than sex-related genes (purple). The y axis shows the tissue specificity using the metric, where 0 is ubiquitous across tissues, and 1 is tissue-specific, and each point is a different gene (see **Supplementary Figure S5** for the individual genes). **(B)** Study proportions of variance in expression resulting from smoking-related autosomal effects are on average higher than that of sex-related autosomal effects. The y-axis shows the proportion of variation. Each point is the proportion of variance explained by that covariate (sex or smoking) in one study, colored by the location of the probes (orange for autosomal, blue for sex chromosomal).

In addition to comparing tissue-specificity, we used variance components analysis (see **Methods 4**) to compare the contributions of sex and smoking to variation in gene expression. We found that, across studies, smoking explains a significantly larger portion of variation in autosomal gene expression than sex (p=0.015), highlighting the importance of considering extrinsic sources of variation in addition to sex (Figure 5B).

### 4. Airway epithelium shows strong patterns of smoking-related differential expression

We first examined the grouped airway epithelium dataset for patterns of smoking and sex-related differential expression. The airway epithelium dataset consists of 444 samples, which is an expanded version of the dataset analyzed by Yang et al (C. X. Yang et al. 2019) (n=211).

We used principal variance components analysis (PVCA) (see **Methods 4**) to examine the overall contributions of the covariates sex, smoking, and a sex-by-smoking interaction effect to variance in expression. Similar to the analysis across tissues, we found that in the Grouped AE study, smoking-related autosomal genes explain a larger fraction of variance than sex-related autosomal genes (see Figure 6A). Additionally, here we see a larger proportion of sex-related variance due sex chromosomal genes versus autosomal genes.

**Figure 6.**
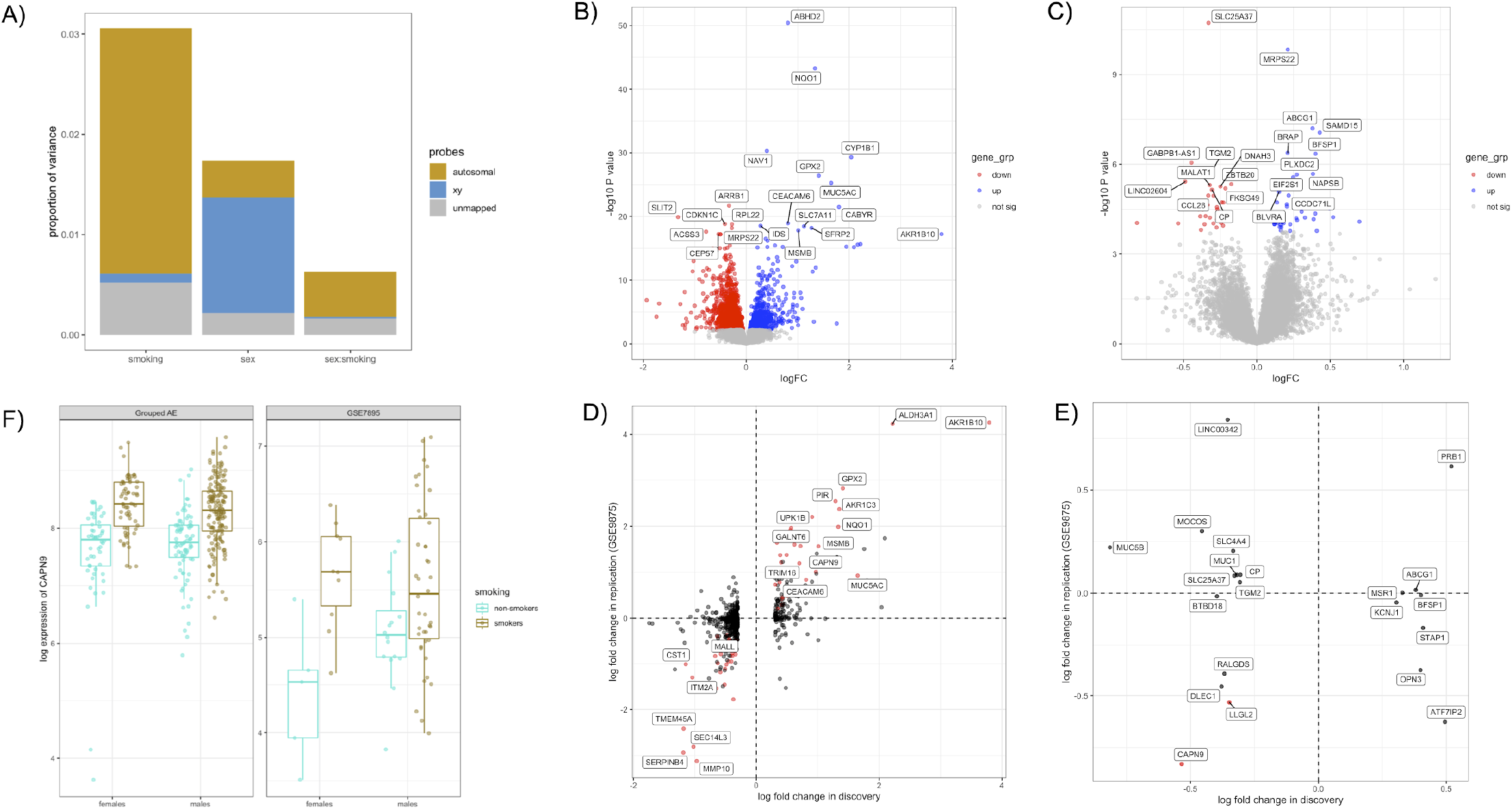
Results from grouped airway epithelium analysis. **(A)** Bar plot showing airway epithelium variance decomposition across smoking, sex, and smoking-by-sex covariates. The location of the probes is given by the color of the bars: orange is autosomal, blue is sex chromosomal, and gray is unmapped. **(B,C)** Volcano plots showing DE genes related to smoking (B) and the sex-by-smoking interaction effect (C). The x-axis is the log-fold change (logFC) in expression between smokers and non-smokers, and the y axis is the −log10 of the unadjusted p-value. Each point is a gene, colored according to significance: red indicates the genes are significantly up in non-smokers, blue indicates the genes are significantly up in smokers, genes in gray do not pass the significance threshold. The top 20 genes (lowest p-value) are labeled. **(D,E)** Replication of DE genes in held out airway epithelium dataset (GSE9875) for sex (D) and sex-differential smoking responses (E). Each point is a DE gene identified in the Grouped AE dataset. The x-axis shows the log fold change in discovery and the y-axis shows the log fold change in the replication dataset. A positive log-fold change corresponds to higher expression in smokers. Red dots indicate genes that pass the replication threshold in the validation dataset. Only the top 20 gene names are shown in (D) for ease of visualization. Dashed lines are at log-fold change zero. **(F)** Visualization of *CAPN9* interaction effects in discovery and validation in female and male smokers (gold) and non-smokers (teal).

We used a model including sex, smoking, and a sex-by-smoking interaction term, in addition to the covariates race-ethnicity, pack-years, age, and submission date. This model is similar to that used by Yang et al. (C. X. Yang et al. 2019) but also includes submission date to account for batch effects (i.e. effect of non-biological factors) seen in the data (see **Supplementary Figure S6**). Using this model with an FDR cutoff of <0.05 and absolute log fold-change cutoff of ≥0.3, we identified 2625 probes differentially expressed related to smoking, 128 related to sex, and 1 related with a significant interaction effect. Given that many probes map to the same gene, we sought to leverage these patterns of multi-mapping by meta-analyzing the values of the probes corresponding to each gene (see **Methods 5-2**). After summarizing probes to genes, the same cutoffs resulting in 932 DE genes related to smoking, 48 genes related to sex, and 30 with sex-differential smoking effects (see **Supplementary Tables S5 A-C**). Of these genes, 43 genes with smoking-related and 33 genes sex-related effects were located on the X or Y chromosomes. Volcano plots showing DE genes related to smoking and sex differential smoking effects are included in Figures 6B and C, respectively. Many of these genes were also identified by Yang et al (C. X. Yang et al. 2019) in their analysis, and show similar effect sizes (see **Supplementary Figure S8** for a comparison of smoking-related genes).

We then sought to assess the extent to which these DE genes were replicated in a held-out airway epithelium dataset. From our list of 21 studies, we selected *GSE7895*, which is the largest airway epithelium dataset (and was also used for replication by Yang et al (C. X. Yang et al. 2019)). This dataset was generated by the same lab as the Grouped AE dataset but was on a different platform and represents a different set of subjects. Figures 6D and E compare the effect sizes in the discovery (Grouped AE) dataset versus the replication (GSE7895) dataset for smoking and sex differential smoking effects respectively. While 110 smoking DE and 18 sex-DE genes replicated (same direction effect size and p-value < 0.05), only 1 of the interaction effect genes replicated: *CAPN9*. *CAPN9* is higher in smokers than non-smokers, but appears to show a slightly stronger effect in females than in males; however, it is important to note that the GSE7895 dataset contains only 5 female non-smokers, so it is difficult to draw conclusions about whether this effect is truly replicated (see Figure 6F).

In addition to examining the replication of particular genes, we also sought to examine the relationship of the effect sizes. Specifically, for DE genes identified in the discovery set, we determined whether the effect sizes in the discovery and validation were related. Between the discovery and validation, while there is a strong correlation in the effect sizes for smoking related effects (Pearson’s =0.63, p< 2*10^-16^), there is no correlation in the effect sizes for sex differential smoking effects (Pearson’s =-0.04, p=0.86). The lack of correlation as well as the single gene in the replication of the sex-differential smoking effects is likely due in part to the small sample size and unbalanced nature of the replication set, but also demonstrates a lack of concordance of effect sizes, even if they are not significant in the replication.

### 5. The majority of smoking-related expression studies are underpowered to detect sex differences in smoking effects

In addition to examining the effects of smoking across tissues, we were interested in assessing whether there are sex-differential responses to smoking. However, large sample sizes are required to have sufficient power to detect interaction effects, which are often very small. Assuming best case scenario where the datasets are balanced - i.e. ¼ each of male smokers, male non-smokers, female smokers, and female smokers - in order to have 80% power to detect absolute log effect sizes of 0.3 (i.e. 1.2-fold difference in expression levels) at an FDR of 0.05, we would need at least 60 samples (see **Supplementary Figure 10** for a visualization of these parameters and **Methods 7** for an explanation of these calculations). It is expected that most interaction effects are smaller than that, and for log effect sizes of 0.2 and 0.1, we would need at least 140 and 525 samples, respectively. The *Grouped AE* study contains 444 samples, but with an uneven breakdown: the smallest category (female non-smokers) contains only 61 samples (14%) and largest (male smokers) contains 200 samples (45%).

The studies overall were highly imbalanced across sex and smoking categories. Across all studies, the median numbers of samples per category are 13.7, 9, 17.3 and 16 samples for female non-smokers, female smokers, male non-smokers, and male smokers, with totals of 424, 279, 535, and 495 samples per category respectively. Only 4 of the 31 smoking-related studies contained at least 15 male and female samples per smoking category (*E-MTAB-3604, GSE17913, E-MTAB-5278, GSE30272)*, and only 2 of these studies have more than 20 males and females per category (*E-MTAB-5278, GSE30272,* with 23 or more per category). The remaining studies did not have sufficient samples for detecting genes with sex-differential smoking effects in standard interaction analyses. Given these power limitations, we focused on whether the interaction effects identified in the *Grouped AE* study replicated in the other studies. None of the 30 genes replicated at Bonferroni corrected p-value threshold (p < 0.05/30). Because this is conservative, we also examined the results at an uncorrected p-value threshold; however, this means that we expect they may be false positives, and all require further validation.

Five of the 30 genes had an uncorrected p-value < 0.05 and same direction effects in the replication: *SLC25A37* and *OPN3* in the study E-MTAB-5278 (blood), and *RALGDS, KCNJ1,* and *MS4A7* in GSE30272 (brain). The list of these genes and their p-values and effect sizes are included in **Supplementary Table 7**; see **Supplementary Figure 11** for visualization of their effects. Briefly, in smokers relative to non-smokers, *SLC25A37* is lower in males and *KCNJ1* is lower in females. Two genes, *OPN3*, *MS4A7,* appear to be lower only in female non-smokers, while *RALGDS* shows opposite direction effects: higher in female smokers and lower in male smokers.

## DISCUSSION

In this study, we sought to examine sex- and smoking-related effects across tissues in publicly available gene expression data. We performed a systematic search of publicly available gene expression datasets, and identified 31 smoking-related studies spanning 1754 samples and 12 tissues as well as an additional group of overlapping airway epithelium studies consisting of 411 samples (which we refer to as the *Grouped Airway Epithelium* study). The studies identified were overall male-biased and unbalanced across smoking and sex-related groups. Only 4 of the 31 studies and the Grouped Airway Epithelium (AE) study contained at least 15 males and females per smoking category.

To our knowledge, our analysis represents the first comprehensive examination of smoking-related gene expression across tissues in publicly available data. Additionally, our analysis concomitantly considers sex-related effects, which are often ignored, and compares the relative impacts of these covariates. We examined smoking-related effects across 31 studies and 12 tissues and found evidence for tissue-specific effects in smoking response, with separate clusters for airway epithelium (and related tissues) and blood. Despite within-tissue similarities, several genes appear to be key players across tissues, including 8 genes (*LRRN3, MS4A6A, GAPDH, RPLP0, CX3CL1, GPR15,* and *AHRR*) that were differentially expressed in both an airway-related and non-airway tissue. Many of these genes have been previously reported to be associated with smoking status. In blood, *LRRN3*, or leucine-rich repeat neuronal 3 gene, has been shown to have increased expression in smokers across multiple studies (Martin et al. 2015; Maas et al. 2020; Huan et al. 2016; Baiju et al. 2021), as well as differential DNA methylation patterns (Guida et al. 2015; Huan et al. 2016). *GPR15* expression is associated with smoking in blood (Huan et al. 2016), *CX3CL1* is associated with lung cancer stage in smokers (Su et al. 2018), and *MS4A6A* is found to have altered DNA methylation in alveolar macrophages in response to smoking (R. A. Philibert et al. 2012). Interestingly, while *GAPDH* and *RPLP0* are housekeeping genes, *GAPDH* has been reported to be differentially expressed in response to smoking in mouse lungs (Agarwal et al. 2012). It is possible that differences in these housekeeping genes highlight differences in numbers and populations of cells, and future work is required to examine potential cell-type specific effects.

By comparison, similar scale sex-related effects appeared to be consistent across studies and tissues. These effects were largely limited to sex chromosomes, which is not unexpected given study size and our use of conservative thresholds. Direct comparison of smoking and sex-related effects highlighted that smoking has a larger impact on autosomal gene expression than sex in the tissues we examined. Many of these tissues were airway-related, so is possible (and likely) that examination of other tissues may show smaller magnitude smoking effects, and we do not know how these effects will compare to sex. Sex-related effects are often overemphasized, and these comparisons illustrate the importance of considering other covariates and disease states that may have larger or similar scale impacts on expression.

In addition to examining overlapping sets of genes and correlations between studies, we used meta-analysis to identify consistently DE genes across tissues, using 27 of the 31 studies as discovery and 4 studies for validation. From this meta-analysis, we identified 7 genes with smoking-related effects: *AHRR, CYP1B1, NQO1, LRRN3* were significantly higher and *ELOVL7*, *CCL4*, and *GZMH* were significantly lower in current smokers as compared to non-smokers (*LRRN3* and *AHRR* were also identified from the study overlap analysis). While the smoking-related genes appeared across studies, only *AHRR* and *GZMH* showed consistent effects across tissues, while the other genes were strongest in a particular tissue: airway epithelium for *NQO1* and *CYP1B1*, blood for *LRRN3,* and alveolar macrophages and sputum for *ELOVL7* and *CCL4.* Four of these genes validated in a held-out set and 4 genes were DE in the validation studies: *LRRN3* (blood - similar tissue specificity), *AHRR* (blood), *NQO1* (airway epithelium - similar tissue specificity), and *CYP1B1* (alveolar macrophages). For sex-related effects, we identified 22 genes, all of which were sex chromosomal and appeared consistent across tissues.

All 7 genes have known associations with smoking. Multiple studies have shown that *LRRN3* is consistently overexpressed in smokers specifically in blood (described above). *NQO1* is overexpressed in airway tissue in response to biofuel smoke (Mondal et al. 2018), matching the possible tissue specificity seen above. However, it has also been shown to be overexpressed in pancreatic tissue of smokers (Lyn-Cook et al. 2006), and a genetic variant located in this gene has an interaction effect with smoking that is associated with colorectal cancer risk (X.-E. Peng et al. 2013). Increased expression of *CYP1B1* in the aerodigestive tract is associated with smoking (Port et al. 2004), and in oral mucosa CYP1B1 has increased expression and differential methylation in smokers vs. non-smokers (Richter et al. 2019). Neither *CCL4* or *ELOVL7* were replicated in our analysis, but have known smoking-related associations. Multiple genetic variants in this *ELOVL7* are associated with smoking behavior (Liu et al. 2019; Wootton et al. 2020) and *CCL4* expression is lowered in PBMCs of smokers (Arimilli et al. 2017).

Multiple studies (Grieshober et al. 2020; Philibert et al. 2020) have found that hypomethylation of *AHRR*, which encodes the Aryl-Hydrocarbon Receptor Repressor, is strongly associated with smoking in several tissues. *AHRR* modulates responses to dioxin toxicity and is involved in regulation of cell growth. Similar to our analysis, additional studies have found that *AHRR* expression is increased in smokers, and decreases following smoking cessation (Bossé et al. 2012). *GZMH* encodes Granzyme H, which is a T and NK cell serine protease involved in lysing target cells. While one study in blood found decreased expression of *GZMH* in smokers (Arimilli et al. 2017), matching our analysis, another study, also in blood, found significantly increased expression (Vink et al. 2017), so further investigation is required to replicate the direction of this effect.

We performed 2 within-tissue meta-analyses for smoking-related effects in blood and airway epithelium, identifying 19 and 66 consistently DE genes, respectively. Interestingly, in airway epithelium, the only overlapping gene, *AKR1C3*, was significantly higher in smokers relative to non-smokers, but in blood, was significantly lower in smokers relative to non-smokers. The significance and direction of effects were replicated in held-out airway epithelium and blood studies, indicating that these opposite-direction effects are robust. To our knowledge, this finding is a novel discovery of a gene that shows opposite-direction, tissue-specific responses to smoking; however, it is unclear why this is the case. Opposite direction effects in different tissues have been reported previously: Obeidat et al.(Obeidat et al. 2017) examined gene expression associations between emphysema in blood and lung, and found that 24 out of 29 overlapping genes showed opposite direction effects across the two tissues. The gene *AKR1C3* encodes an aldo/keto reductase, which is a family of proteins known to be involved in cancers, including head and neck, bladder, prostate, uterine, breast, and ovarian cancer. Other members of the *AKR1C* family are known to be upregulated in response to smoking (Woo et al. 2017), and were similarly found differentially expressed in multiple tissues in our analysis. Examination of *AKR1C3* regulation and tissue-specific expression of genes in nearby pathways may help elucidate this differential response.

For the Grouped AE study, we found 932 significantly DE genes with smoking-related effects, 48 DE genes related to sex, and 30 genes with sex-differential responses to smoking. This is an expanded re-analysis of the samples examined by Yang et al. (C. X. Yang et al. 2019) (n= 211 samples). Despite our larger sample size, we identified fewer genes because we used more conservative thresholds and included an additional batch-related covariate. There was both substantial overlap and correlation between effect sizes for the smoking-related effects, but not for the sex-differential smoking effects. It is possible that we did not observe a correlation for the sex-differential smoking effects because the replication study was very small. Additionally, while 110 smoking DE genes and 18 sex DE genes replicated, only 1 gene with a sex differential smoking effect, *CAPN9*, was replicated in the validation study. Both male and female smokers showed increased expression of *CAPN9*, but this increase appears to be slightly stronger in females relative to males; however, this effect is subtle and the replication dataset was unbalanced, with only 5 non-smoking females. *CAPN9* encodes a calcium-dependent cysteine protease, which is activated in response to oxidative stress, and its expression is inversely associated with prognosis in gastric cancer (P. Peng et al. 2016). Additionally, a previous study found that *CAPN9* was correlated with the expression of *MUC5AC*, which is a mucin gene known to respond to smoking (Goldfarbmuren et al., n.d.).

We found that the majority of the remaining publicly available smoking studies were too small to identify sex-differential smoking (or sex-by-smoking) effects on gene expression. Additionally, most studies were unbalanced, decreasing power to detect these effects. Only 4 studies had at least 15 samples per sex/smoking category, with a maximum of 23 samples in the largest of these 2 studies. Due to the limited sample sizes, we used these studies to examine replication of the 30 sex-differential smoking genes identified in Grouped AE. No genes were replicated after correcting for the number of tests (n=30). At a nominal p-value cutoff (uncorrected p < 0.05), 5 genes were identified that showed the same patterns in the discovery and validation: *SLC25A37* and *OPN3* in the blood study and *RALGDS, KCNJ1,* and *MS4A7* in the brain study. It is important to note that the studies were from various tissues (blood, brain, sputum, and buccal mucosa) and not airway epithelium, so it is possible that the lack of replication was in part due to tissue specificity; however, it may be due to sample size. We cannot draw conclusions about replicability or tissue-specificity of sex-related smoking effects without examining larger validation studies.

This work has several strengths. First, we performed a systematic search to identify and manually filter smoking-related studies available in public gene expression databases in order to construct our compendia of smoking studies. By performing such a search, we ensured that we obtained a comprehensive picture of smoking effects on gene expression, rather than cherry-picking specific studies. We also leveraged our previously developed method (Flynn, Chang, and Altman 2021) to infer sex labels for these studies, without which, 9 of the 31 studies would not have been available for analysis. As part of this sex labeling process, we also discarded samples with mismatched metadata and inferred labels, which may also have other mislabeled metadata, thereby increasing the quality of our data.

In our analysis of smoking and sex-related effects, we made conservative methodological choices in order to identify consistent, reproducible effects. Our cutoff for identifying DE genes consisted of both an effect size and FDR threshold. Additionally, we employed meta-analytic techniques to summarize probes to genes in our comparisons, which has been suggested before in the literature (Ramasamy et al. 2008), but to our knowledge not yet employed. We demonstrate that use of this technique decreases the number of false positives. It is important to note that meta-analysis also increases bias toward genes with more probes, which is a concern for consistent examination across genes; however, it does not present problems if concerned with true positive rate. By making these choices, we expect that our analysis has false negatives and that we may have missed some subtle effects.

Two additional strengths of our analysis are the examination of the correlation structure between studies and the side-by-side comparison of smoking and sex-related effects. Using a weighted correlation metric allowed us to better understand the overall pattern of replication without relying on specific significance cutoffs, which both require making decisions about a threshold and could potentially miss replicated genes because of small sample sizes. The concurrent analyses of smoking and sex-related effects allowed us to compare the tissue specificity of the two effects. Sex-related gene expression has been examined across tissues extensively (Gershoni and Pietrokovski 2017; Oliva et al. 2020; Mayne et al. 2016), and has been shown to have both strong, shared sex chromosomal effects and small tissue-specific autosomal effects. In our analysis, in part because of sample size and effect size cutoffs, we only saw sex chromosomal effects which were present across tissues. This is in contrast to the smoking-related effects that showed some tissue-specific patterns, which we identified in the same studies at the same significance thresholds.

While the use of public data is a strength of our analysis, it also presents a limitation. Larger studies on which previous analyses have been performed (Bossé et al. 2012; Huan et al. 2016; Maas et al. 2020) are either not publicly available or missing sufficient metadata for re-analysis of sex-related effects. Public data is also biased toward specific tissues, and while we sought to examine effects across tissues, we were limited to the seven tissues with data available. The majority of the available tissues were airway-related or blood, which makes sense given the nature of smoking-related exposures and ease of sampling peripheral blood, but does not provide a complete picture. Additionally, with the exception of airway epithelium and blood, which had at least 5 studies each, there were less than 3 studies per tissue and many tissues with only 1 study (e.g., brain, liver, kidney), which prevented an assessment of the extent to which some smoking-related effects are tissue (rather than study) specific. Much of the data were also generated by a single lab and on similar platforms. While this lack of heterogeneity makes the analysis less complex, increased heterogeneity in studies leads to identification of more robust, reproducible effects.

We also relied on the author-processed expression data for each study, which helped us obtain data from a heterogeneous set of platforms. However, different processing pipelines are known to greatly affect microarray results (Ioannidis et al. 2009). These effects are disproportionately on the sex chromosomes (Castagné et al. 2011), which may have led to an underestimation of sex chromosome contributions to variance. This also limited our analysis to studies with available processed data. Use of standardized processing steps will allow us to examine additional studies, and may reduce heterogeneity between studies due to processing artifacts. We also limited our analysis to samples from healthy tissues; however, future analyses may include disease samples, which may increase the search space and enable examination of additional questions. In the process of identifying the studies for our analysis, we also identified 47 studies that involved cultured cells exposed to smoke components. While it is unclear whether sex-related effects identified in culture would translate to humans, use of these data, which have many replicates and show larger magnitude smoking responses could help identify sex-related smoking effects.

Many studies were missing important covariate information, including age, race/ethnicity, pack-years, and batch-related effects. Available covariates were included in our models; however, this may have led to inconsistencies across studies because of differing sets of covariates. For studies with missing covariate information, confounding may contribute to the identified genes, leading to incorrect associations. For example, because men smoke more heavily on average (Baumert et al. 2010), without pack-years information, effects attributable to smoking amount might be attributed to sex. In addition to variation in available covariates, studies have shown that self-reported data on smoking is often inaccurate (Gorber et al. 2009). Some studies use plasma or urine cotinine levels to confirm smoking status; however, only 1 study reported these levels. As a result, definitions of smoking may be inconsistent across studies and may include incorrect labels due to self-report or sample label mix ups (while our sex labeling method detects samples with swapped sex labels, we cannot detect mislabeling if it occurs between samples of the same sex). Future work may involve developing models to infer additional covariates and detection of mislabeled samples in other dimensions, such as for smoking status. A possible direction could involve training models to infer smoking status from expression data using either previously identified tissue-specific gene signatures (e.g. (Bossé et al. 2012; Martin et al. 2015)) and/or genes identified in our meta-analysis. This could allow us to expand our analysis to many additional studies that do not contain smoking metadata.

Another limitation is that our study focuses on gene expression data: smoking-related effects occur on multiple biological levels, some of which have sex-related differences. In tumor microenvironments, changes in immune cell populations in response to smoking were more pronounced in women than in men (Alisoltani et al., n.d.). DNA methylation shows sex-specific changes in response to smoking (Koo et al. 2020). Examination of these molecular data types in concert with expression data may help identify additional important insights into smoking and sex-related smoking effects.

In conclusion, we performed a large-scale systematic analysis of smoking and sex-related smoking effects in healthy participants using publicly available gene expression samples from 31 studies and 1 study compendium, spanning 12 tissues. This analysis is the first to examine these effects at this scale and in a sex-aware manner. Our results indicate that expression changes in response to smoking largely cluster by tissue while also showing consistent effects across tissues in a small number of genes. This is in contrast to similar magnitude sex-related effects, which appear to be consistent across tissues. Comparison of smoking and sex-related effects indicate that smoking has a larger impact on autosomal expression than sex in the tissues examined in this study. Our study also highlights the challenges of examining and replicating sex-differential smoking effects in publicly available data, which is in part due to sample size and sex bias. Expansion of this analysis to additional studies and samples may help to validate and further examine patterns of tissue-specificity and assess sex-differential smoking effects.

## Acknowledgements

This work was supported in part by a Center of Excellence in Regulatory Science and Innovation (CERSI) grant to University of California, San Francisco (UCSF) and Stanford University from the US Food and Drug Administration (U01FD005978) Office of Women’s Health. Its contents are solely the responsibility of the authors and do not represent the official views of HHS or the FDA. The authors would like to thank Drs. Jonathan Kwan and Carolyn Dresler for their input on study design.

The majority of the computing for this project was performed on the Stanford University Sherlock cluster. We would like to thank the Stanford Research Computing Center for providing the computational resources that contributed to these research results.

## Author contributions

EF, AC, BN, and RBA conceived the study together. EF and AC performed the systematic search for studies and data processing together. EF performed most downstream analyses and data visualization, and AC assisted with this and on interpretation of the results. BN and RBA supervised the project and provided regular feedback. All authors contributed to writing the manuscript.

## Data and code availability

All data used in this analysis is freely available on GEO or ArrayExpress: study accessions for are located in **Supplementary Table 1** (for grouped airway epithelium) and Table 1B (for all other tissues). The code used in the analysis is available on github at: https://github.com/erflynn/smoking_sex_expression.

