## Supplementary Figures and Captions for "Comprehensive Assessment of Smoking and Sex Related Effects in Publicly Available Gene Expression Data"

**Supplementary Tables and Figures for:**
*Comprehensive Assessment of Smoking and Sex Related Effects in Publicly Available Gene Expression Data*

Emily Flynn, Bridget Nugent, Annie Chang, Jonathan Kwan, Carolyn Dressler, and Russ Altman

Supplementary Table Captions

**Supplementary Table 1.** List of studies included in the Grouped Airway Epithelium dataset with their corresponding PubMed IDs, authors, titles, and citations.

**Supplementary Table 2.** Sex breakdown of samples and studies for *metadata* and *expression*-based labels, along with numbers of *unlabeled* samples. The fields for numbers of *mixed sex*, *mostly-male*, and *mostly-female* are left blank for samples because they are only defined for studies (see **Methods 2** for definitions).

**Supplementary Table 3.** Summary of results across studies for smoking and sex related effects. For each of the studies, the estimated *proportions of variance* (see **Methods 4**) are shown for sex and smoking as well as for sex-by-smoking interaction effects, along with the numbers of *differentially expressed* probes and genes for sex and smoking. In addition, the *total* number of *probes* and *genes* for each study's platform is listed. Sex-related effects are not included for the three single-sex studies.

**Supplementary Table 4.** Lists of DE genes present in at least two (non-Grouped AE) studies for smoking (A) and sex (B). For each gene, the *chromosome* location, and the number of studies (*nstudies*) as well as the study names (*studies*) and *tissues* are listed, along with the group the gene is more highly expressed (*overexpressed*) in: either smokers or non-smokers, or males or females.

**Supplementary Table 5.** Results for sex and smoking meta-analyses across tissues (*all*) and for *blood* and *airway* epithelium individually. Genes resulting from the meta-analysis are listed with their *chromosome* location and number of studies (*nstudies)* they are present in. The identities of validation studies (*valid.study*) are included for genes that are validated in held-out data. Log-fold change (*logFC*), standard deviation (*SD*), and p-values (*p)* are included for both the discovery and validation. The FDR-adjusted p-value (*adj.p*) is also provided for the discovery.

**Supplementary Table 6.** DE genes in Grouped Airway Epithelium for (A) smoking, (B) sex, and (C) sex-by-smoking. For each gene, its *chromosome* location*,* log-fold change (*logFC),* standard deviation (*SD*), p-value (*p)*, and FDR-adjusted p-value (*adj.p*), as well as the number of probes (*nprobes*) present for that gene in the array.

**Supplementary Table 7.** Sex-differential smoking effects in grouped AE replicated at a nominal p-value. For each gene, the name of the validation study (*valid.study*), along with discovery and validation log-fold-changes (*logFC*) and p-values are included. For the discovery, the FDR-adjusted p-value is listed (*adj.p.disc*).

### Supplementary Figures

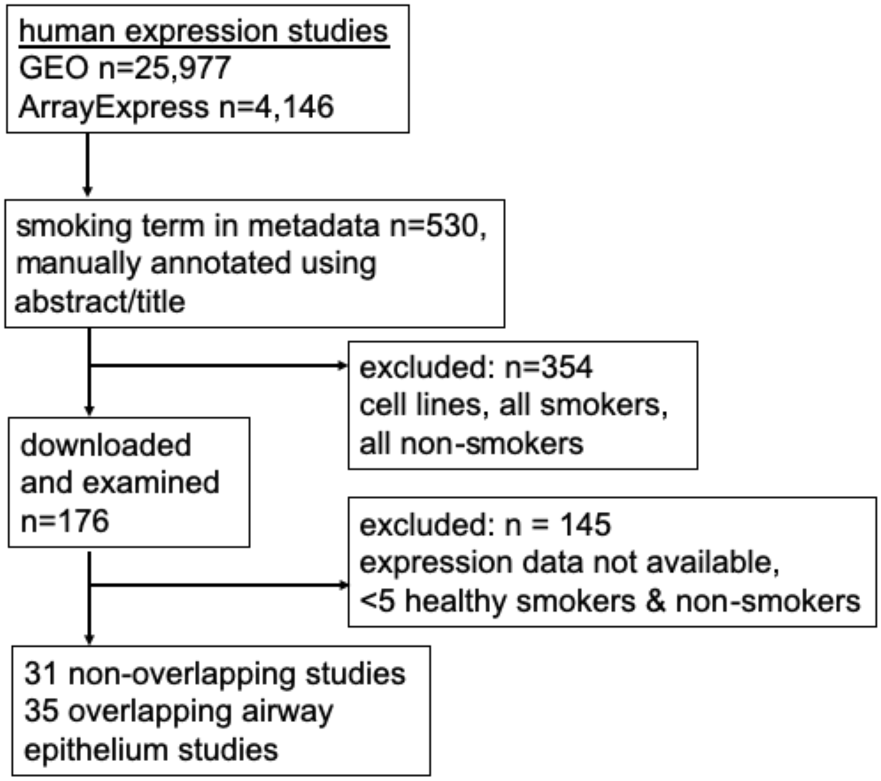


**Supplementary Figure 1.** Diagram of systematic search for smoking-related data.

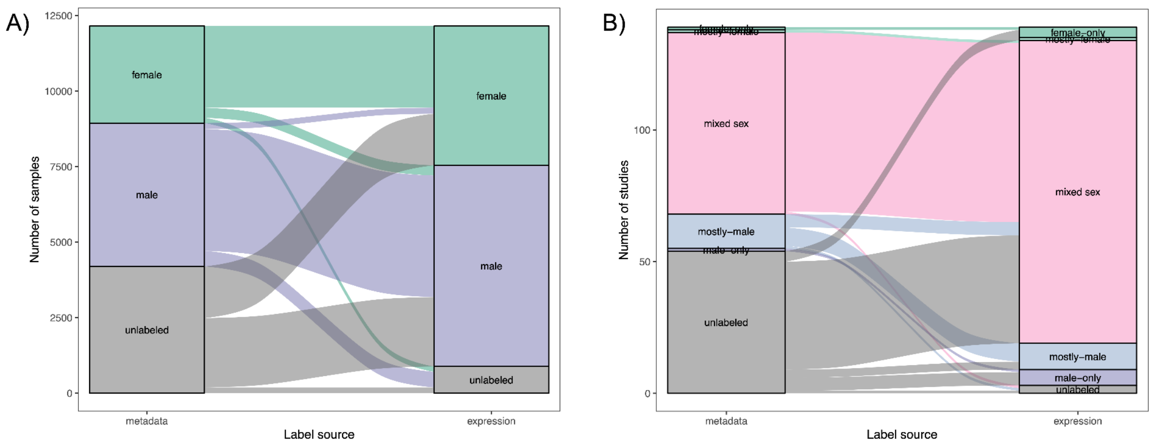


**Supplementary Figure 2**. Alluvial diagram showing bias in smoking data before and after inferring missing sex labels in samples (A) and studies (B) in smoking history. In each panel, the left-hand column includes the metadata labels while the right-hand column is the expression inferred sex labels, each color represents a different label: green is female, blue is male, and gray is unlabeled. For B) pink indicates mixed sex studies, and pale blue indicates mostly-male studies. See **Supplementary Table 2** for the corresponding values.

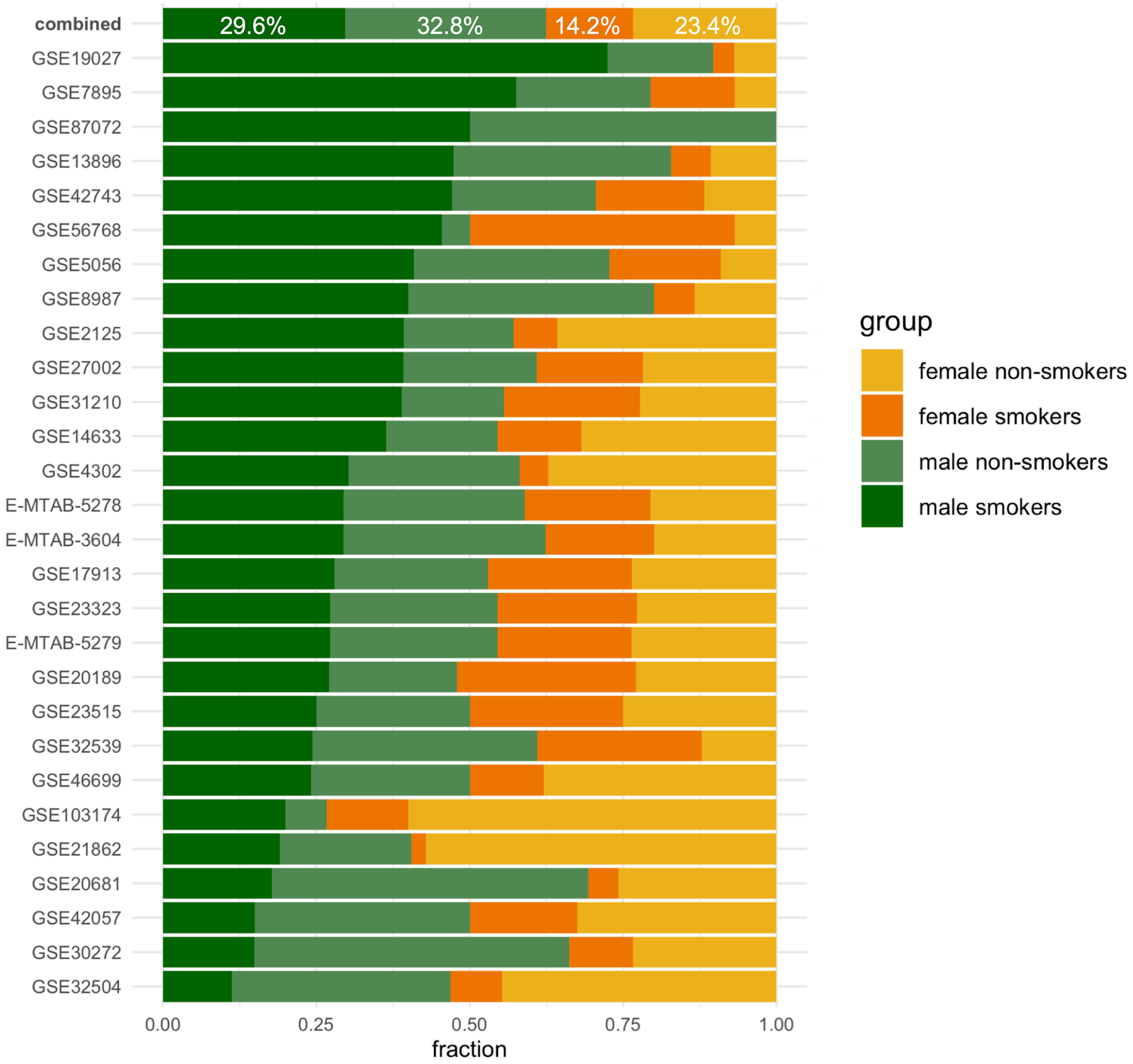


**Supplementary Figure 3.** Sex and smoking breakdown for the 28 mixed sex studies (1588 samples) included in the analysis. Each color represents the fraction of the population in each sub-group: *green* indicates males and *orange* indicates females, the *darker* versions of each color are used for smokers. The combined breakdown is shown at the top with percentages listed for each fraction. For counts of the number of samples per study, see **Table 1B**.

A)

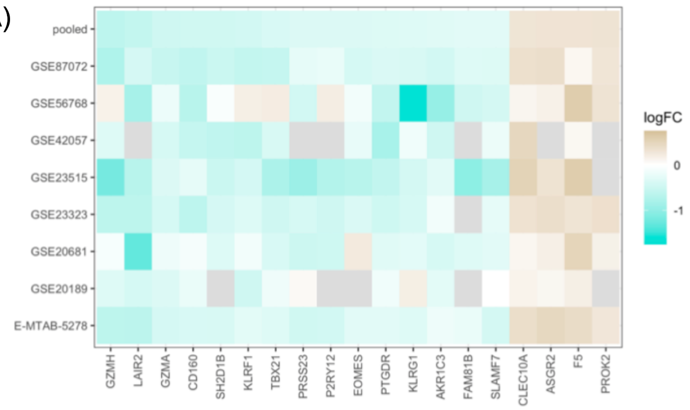


B)

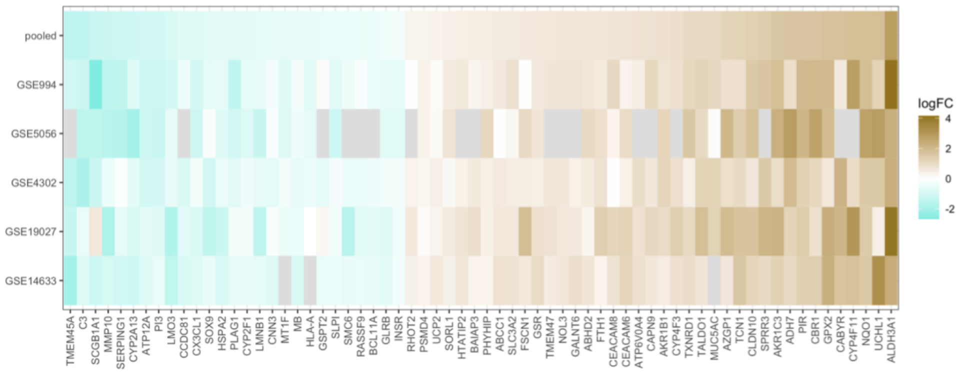


**Supplementary Figure 4.** Meta-analysis of smoking DE genes in blood (A) and airway epithelium (B). Gold is upregulated and teal is downregulated in smokers vs non-smokers. Gray tiles indicate missing values. The resulting genes are also listed in **Supplementary Table 5**.

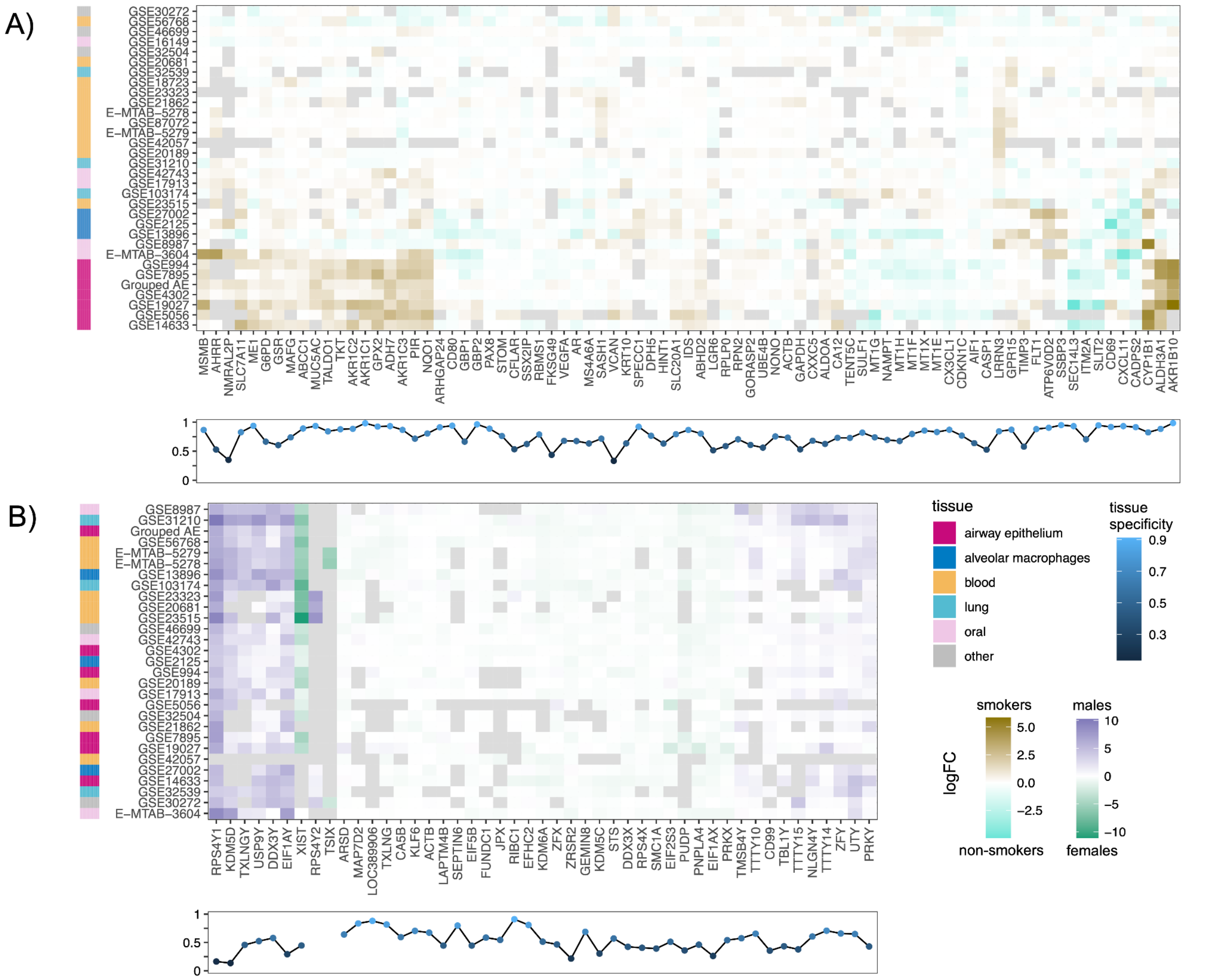


**Supplementary Figure 5.** Examination of tissue-specificity among smoking **(A)** and sex **(B)** DE genes. Heatmaps are shown for genes DE in at least 2 tissues and 3 studies. Heatmaps were clustered by both rows and colors. The bar along the left side of the heatmaps indicates the study tissue (for ease of simplicity and ease of visualization, tissue categories were manually collapsed into larger groups, including blood, oral, and other). Heatmap colors are as follows: turquoise indicates upregulation in non-smokers and blue is upregulation in smokers, purple is upregulation in males and green is upregulation in females, and gray indicates missing values. Underneath the heatmaps, the tissue specificity metric ($\tau$, see **Methods 5-4**) is shown for each gene, both plotted from 0 to 1 and also colored based on this value, with dark blue (or values toward 0) indicating ubiquitous expression across tissues and light blue (or values toward 1) indicating high tissue-specificity.

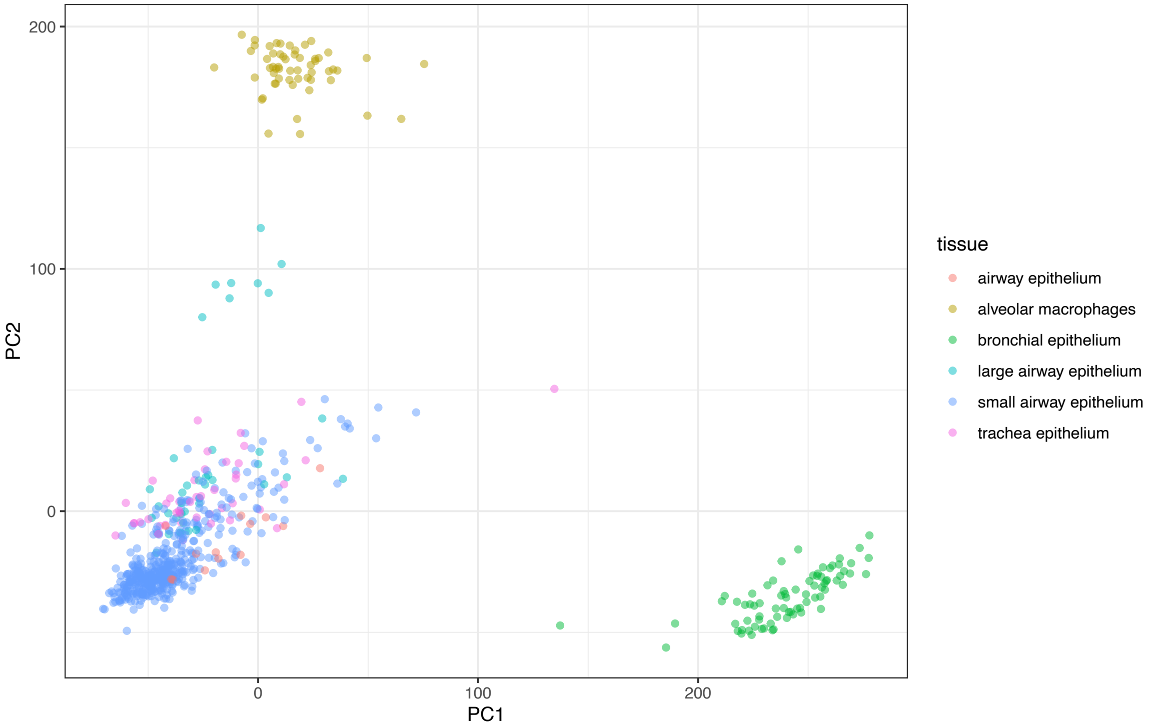

PC2
PC1
200
100
0
100
200
tissue
airway epithelium
alveolar macrophages
bronchial epithelium
large airway epithelium
small airway epithelium
trachea epithelium


A)

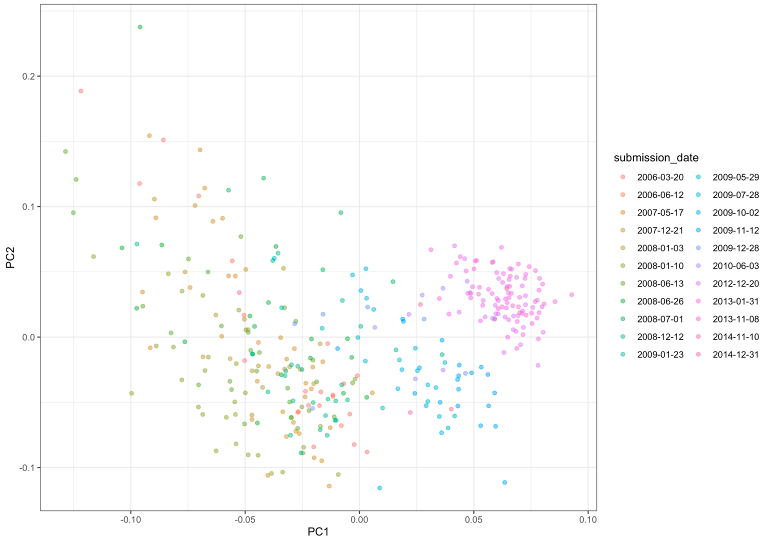

PC2
PC1
0.2
0.1
0.0
-0.1
-0.10
-0.05
0.00
0.05
0.10
submission_date
2006-03-20
2006-06-12
2007-05-17
2007-12-21
2008-01-03
2008-01-10
2008-06-13
2008-06-26
2008-07-01
2008-12-12
2009-01-23
2009-05-29
2009-07-28
2009-10-02
2009-11-12
2009-12-28
2010-06-03
2012-12-20
2013-01-31
2013-11-08
2014-11-10
2014-12-31


B)

C)

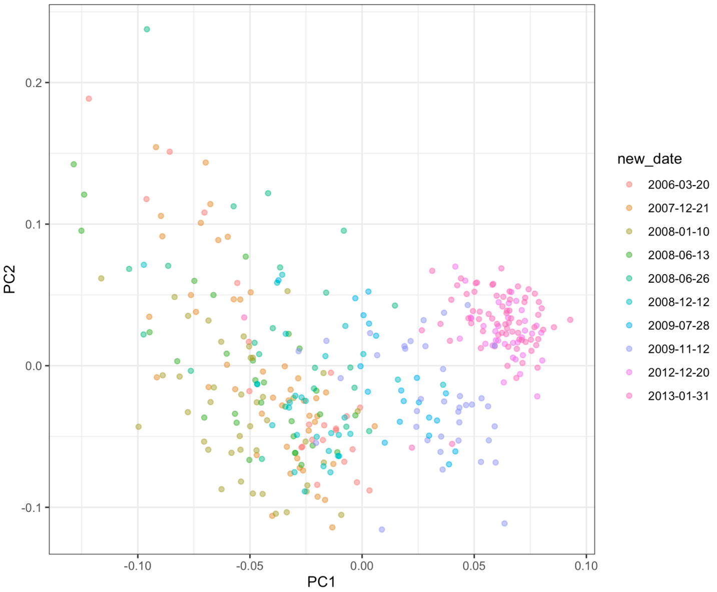


**Supplementary Figure 6.** PC space visualizations. A) Small and large airway epithelium cluster together in principal component space. B) Airway epithelium colored by submission date, showing batch effects, and C) Colored by date after collapsing small dates.

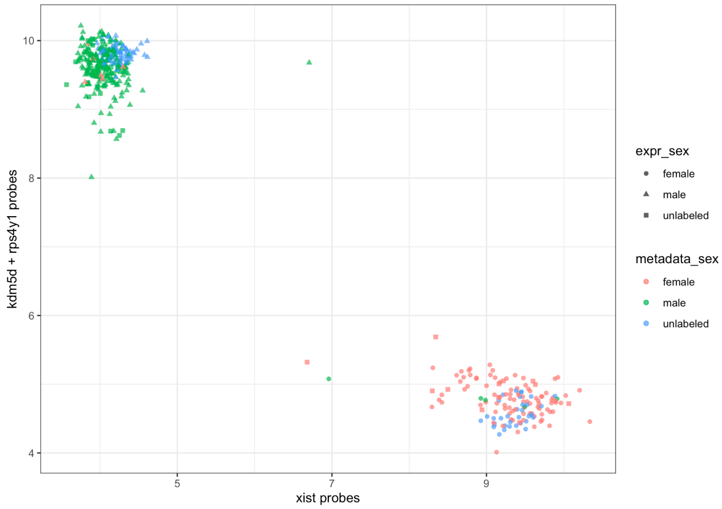


**Supplementary Figure 7.** Clustering of sex labels for airway epithelium data using the Toker method (Toker et al., 2016), which relies on the expression of XIST probes, which are only expressed in females, and RPS4Y1 and KDM5D probes which are only expressed in males. The averages of these sets of probes are shown on the X and Y axis respectively. Each point represents a sample, samples are colored by metadata sex (red=female, green=male, blue=unlabeled) and the shape relates to the expression-based inferred sex (circle=female, triangle=male, square=unlabeled).

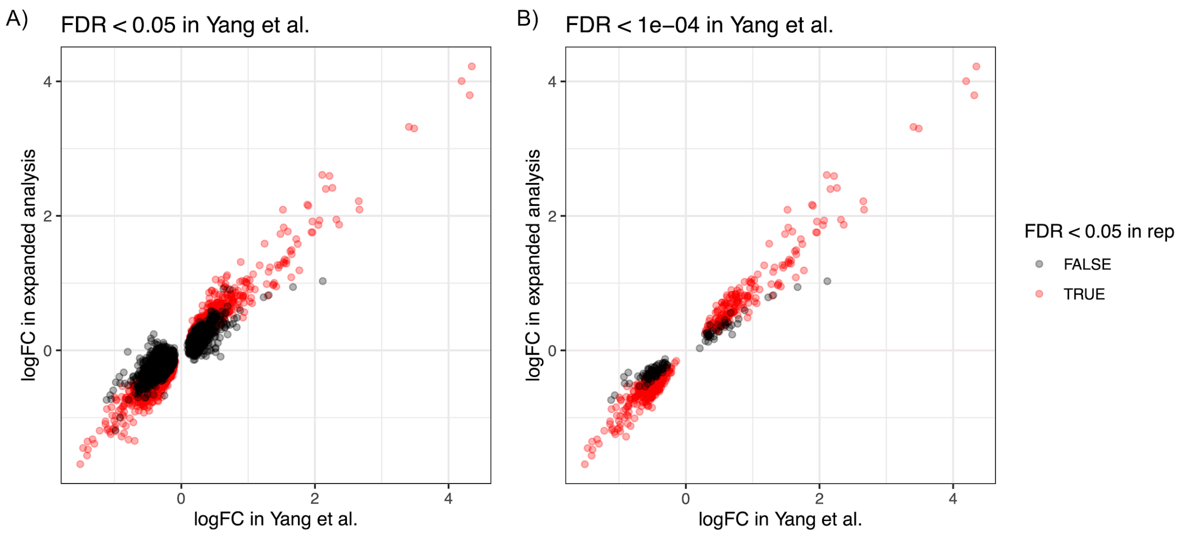


**Supplementary Figure 8.** Comparison of the effect sizes of smoking-related effects between our expanded analysis of the Grouped Airway Epithelium dataset (n=444 samples), and the previous analysis performed by Yang et al. (C. X. Yang et al. 2019) (n=211). Each point is a DE gene, and the genes are colored by whether or not they were replicated in Yang et al.'s analysis (red indicates replicated, black indicates not replicated). The plots show sets of DE genes from Yang et al. at 2 different FDR cutoffs: 0.05 **(A)** and 0.0001 **(B)**. As expected, we see greater overlap and higher numbers of replicated genes at the more stringent threshold.

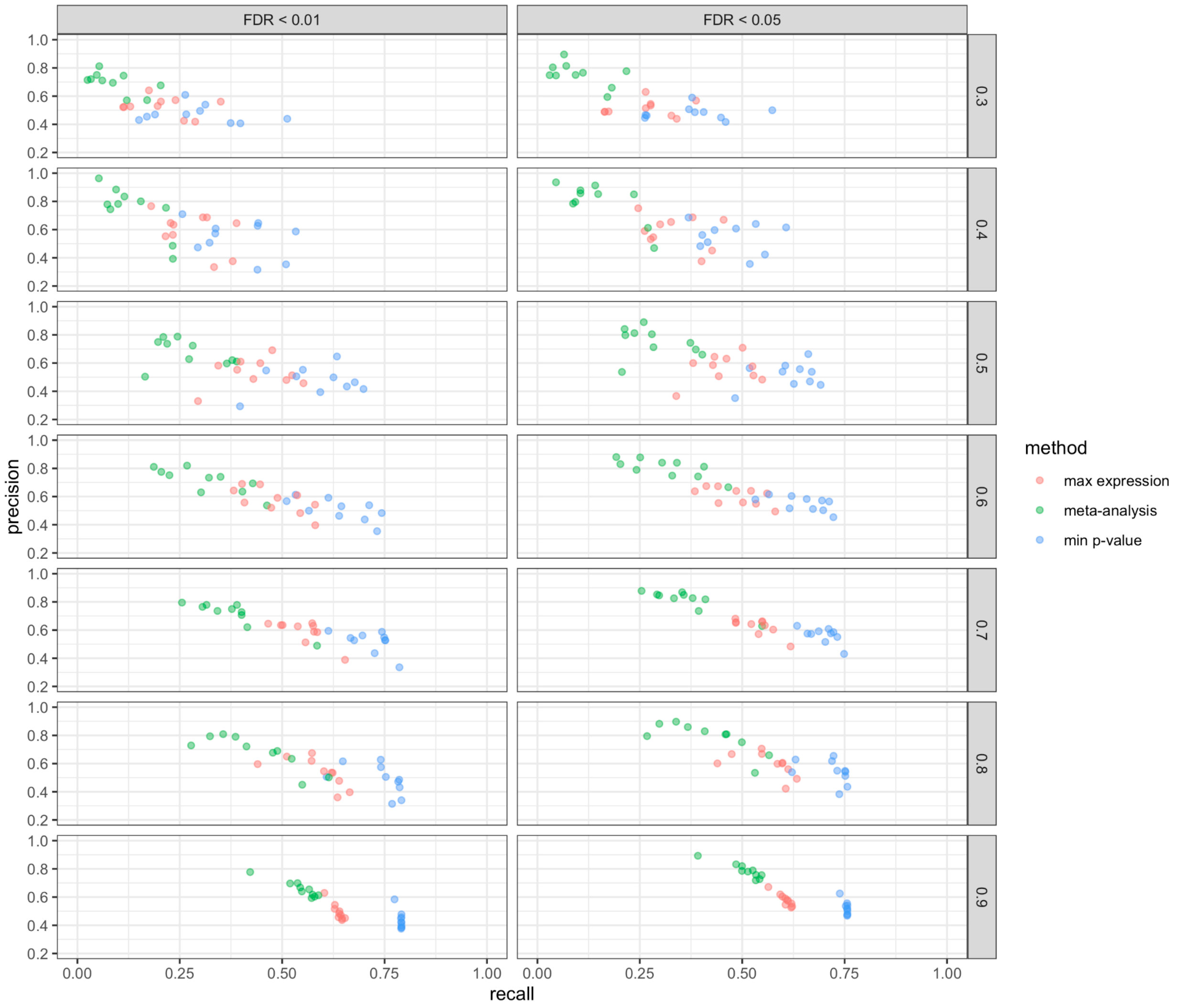


**Supplementary Figure 9.** Comparison of probe-to-gene summarization methods. We compared summarization of probes-to-genes using 3 methods: (1) fixed effects meta-analysis (green), (2) selecting the probe with the highest expression (red), and (3) selecting the probe with lowest p-value (blue). Because there is no ground truth, we assessed the ability of subsets of a dataset to identify DE genes in the entire dataset. Each row panel is a different fraction of the full dataset, and each column panel is the FDR cutoff. Precision and recall are calculated by counting numbers of genes found in that dataset fraction using the method and their overlap with the entire dataset and are shown on the x and y axes respectively. Each point is the result of a randomized run, 15 of which were performed for each dataset fraction.

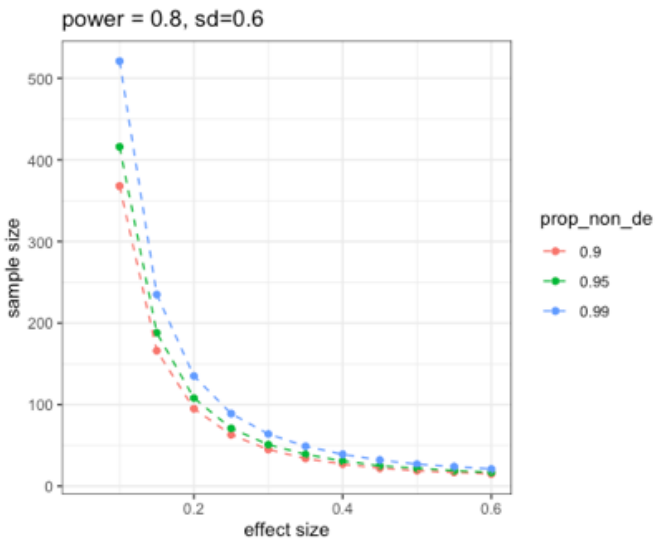


**Supplementary Figure 10.** Sample size calculation for binary interaction effects in expression data. The x-axis is the effect size in log-scale and the y axis is the sample size of the study. Here we assume that the sample is balanced among all four categories. Calculations are shown at power=0.80 and FDR=0.05 assuming a standard deviation of 0.6 across genes. The color of the points shows the results under the assumption that we expect 90%, 95%, and 99% of genes to be not differentially expressed (see **Methods 7** for more information).

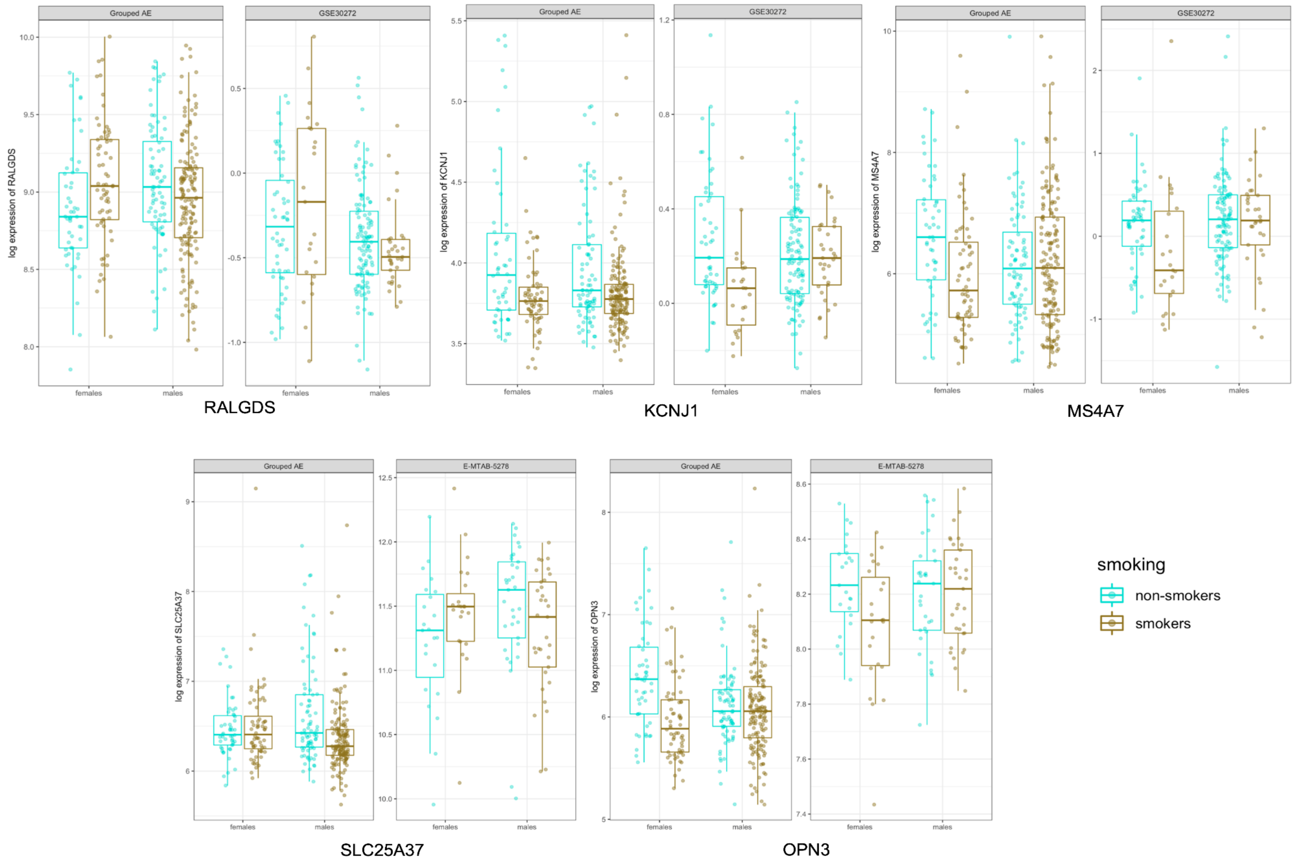


**Supplementary Figure 11.** Five genes that show a nominal interaction effect in replication data. For each gene, the first panel are the results from Grouped Airway Epithelium and the second panel is the replication study (GSE30272 in the first row, and E-MTAB-5278 in the second row). The y axis is the log expression of the gene (scaled for each panel). The effects are shown in female and male non-smokers (turquoise) smokers (gold). Each point is the expression in an individual sample.
